# Bidirectional Electron Transfer in Far-Red Light Adapted Photosystem I: Implications for the Photosystem’s Functionality

**DOI:** 10.64898/2026.07.21.739882

**Authors:** Andrea Calcinoni, Anna Paola Casazza, Alessandro Agostini, Marco Bortolus, Donatella Carbonera, Stefano Santabarbara

## Abstract

Far-Red (FR) Light Photoacclimation (FaRLiP) enables cyanobacteria to extend photosynthetic activity into the far-red region by extensively remodelling Photosystem I (PSI), including the replacement of several core subunits with paralogs that coordinate the red-shifted chlorophyll *f* (Chl *f*). The binding positions of Chls *f* are still a matter of debate, with the most recent structural findings supporting the location of a single Chl *f* molecule within the reaction centre (RC) at the so-called A_-1B_ site. This was in turn suggested to strongly affect electron transfer (ET) directionality leading to an almost monodirectional transfer along the B branch in FR-PSI RC. Here, we directly probe ET in FR-PSI by characterising the photogenerated [P_700_⁺A_1_^−^] spin-correlated radical pair using complementary pulse and Time-Resolved (TR) Electron Paramagnetic Resonance (EPR) spectroscopy at cryogenic temperature. Electron spin-echo decay kinetics are distinctly biexponential, indicating the formation of two charge-separated states. Consistently, out-of-phase ESEEM traces are quantitatively described by two modulation frequencies arising from different dipolar interactions, while TR-EPR spectra are accurately simulated by the combined contributions of [P_700_⁺A_1A_⁻] and [P_700_⁺A_1B_⁻] radical pairs. These results provide direct spectroscopic evidence that both the A and B branches remain photochemically active in FR-PSI. The conservation of bidirectional ET, even when considering the presence of a single Chl *f* molecule in the RC, further implies that the two radical pairs originate from a common primary electron donor. This finding identifies P_700_ as the most likely primary donor and argues against a mechanism in which the RC Chl *f* initiates charge separation.

## 1. Introduction

Far-Red Light Photoacclimation (FaRLiP) is a complex and reversible adaptive response that enables certain cyanobacterial species exposed to light highly enriched in the far-red (FR) region of the solar spectrum to extensively remodel their photosynthetic apparatus, thereby sustaining growth under these otherwise limiting conditions.^1–3^ The FaRLiP response involves the expression of specific protein isoforms of both Photosystem I and II (PSI, PSII), including the main cofactor-binding subunits, together with paralog subunits of the phycobilisomes, accompanied by a profound reorganisation of the supramolecular architecture of the photosynthetic apparatus.^1,3–5^ Most importantly, FaRLiP cyanobacteria synthesise Chlorophylls (Chls) *d* and *f*,^1,3,6^ whose absorption and fluorescence maxima are substantially red-shifted relative to those of the canonical Chl *a*.^7,8^ These intrinsically far-red-shifted pigments are selectively incorporated into PSII and PSI through the expression of Chl *d/f*-compatible protein isoforms,^9^ thereby extending the spectral range of light harvesting into the far-red region and enabling photochemistry under low-energy illumination. Efficient photosynthesis under far-red light, however, requires precise tuning of excitation energy transfer between antenna pigments and the reaction centres (RCs) of the photosystems, as the presence of low-energy chlorophylls creates energetic sinks that may compete with the RCs for excitation trapping, potentially slowing or impairing the charge separation reactions that initiate photochemistry. Consequently, identifying the locations and photophysical properties of Chls *d* and *f*, and, more importantly, elucidating how these pigments reshape excitation energy flow toward the RCs, has remained a central challenge that motivated extensive experimental and theoretical investigations.

With regard to FR-PSII, a general consensus has emerged that one of the RC Chl *a* molecules is replaced by Chl *d* at the so-called Chl_D1_ position^10–12^ where it has been proposed to participate directly in primary charge separation.^13–16^ The incorporation of a single far-red absorbing Chl inside FR-PSII RC is regarded as a key adaptive feature that enables efficient charge separation under FR light while simultaneously compensating for the presence of low energy Chl *f* molecules in the antenna by preserving favourable energetic connectivity to the RC.

In contrast to FR-PSII, the functionality of PSI after acclimation to FR light remains controversial. Whereas there is unanimous agreement that Chl *f* participates in light harvesting, its direct involvement in photochemical reactions is still actively debated. Evidence in favour of Chl *f* replacing Chl *a* in the RC emerged from the light-induced oxidised-*minus*-reduced absorption spectrum, displaying a distinctive derivative-shaped feature at ∼755 nm that was interpreted as an electro-chromic shift of Chl *f* molecules, induced by the positive charge on P_700_^+^, suggesting their location at the positions commonly labelled A_-1A/B_ (structural sites eC-B2/eC-A2 respectively).^10^ The interpretation was also supported by spectroscopic works based on difference-IR spectroscopy.^17,18^ Early ultrafast visible transient absorption measurements also considered the possibility that Chl *f* is incorporated into the electron transfer (ET) chain, although these studies could not conclusively distinguish this scenario from one in which Chl *f* operates exclusively as a light-harvesting pigment.^19^ On the other hand, later femtosecond transient absorption measurements on isolated FR-PSI excited with FR pulses revealed slower formation of charge-separated states than in canonical white-light PSI,^20^ an observation attributed to uphill excitation energy transfer from low-energy Chl *f* antenna states to an unchanged RC, thereby arguing against the participation of Chl *f* in the ET chain.^20,21^ Substantial structural support for this model has been provided by cryo-electron microscopy studies of FR-PSI complexes from multiple FaRLiP cyanobacteria, which identified putative Chl *f* binding sites at peripheral positions within the FaRLiP-specific PsaA and PsaB subunits, while failing to detect Chl *f* molecules within the RC itself.^22–25^

Yet, a very recent high-resolution cryo-electron microscopy study claiming to provide a comprehensive assignment of all Chl *f* binding sites in the FR-PSI of *Chroococcidiopsis thermalis* PCC 7203,^26^ proposed the replacement of Chl *a* at position eC-A2 (A_-1B_), on the B-branch of the ET chain, with Chl *f* (**Figure 1**).^26^ Such single Chl *a*-to-*f* substitution would facilitate the excited state localisation on the RC, making this Chl *f* part of the primary photochemistry events.^26^ Introducing this large asymmetry in the FR-PSI RC composition (and thus energetics) is predicted to strongly favour photochemistry and successive ET along the PsaB-cofactor branch. This scenario represents a significant difference with respect to canonical Chl *a*-only binding PSI, in which the occurrence of bidirectional ET along the cofactor chains coordinated by both the PsaA and PsaB subunits is widely accepted as the dominant mechanism.^27–29^

**Figure 1.**
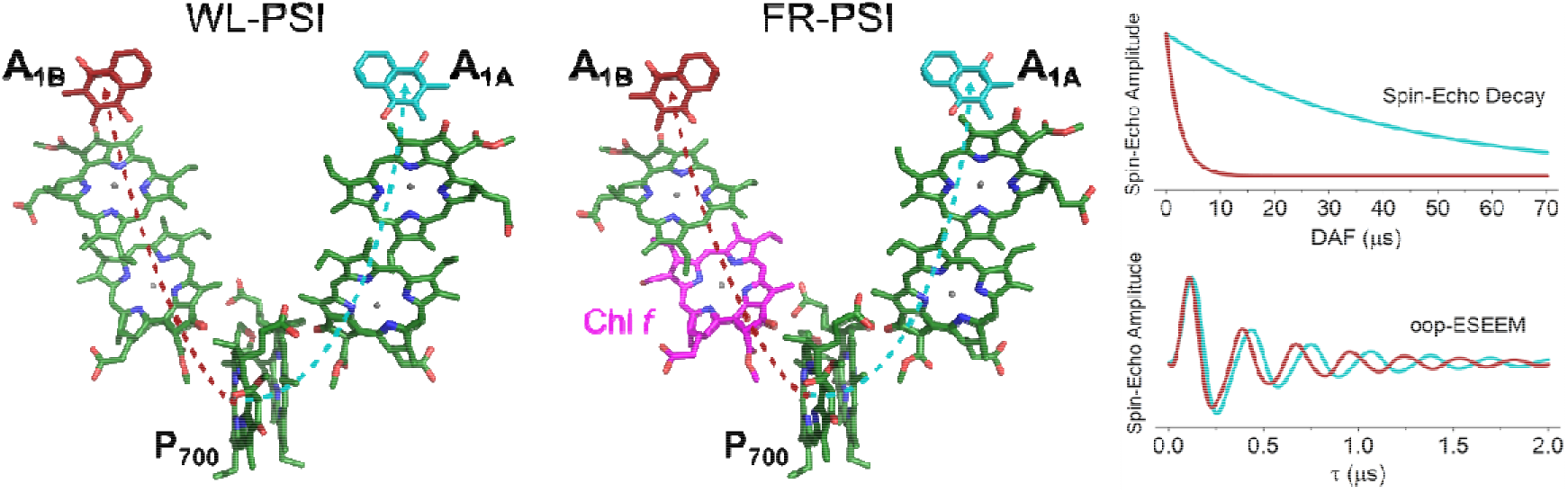
Cofactor arrangements in WL- and FR-PSI RC, along with the expected spin-echo decay and oop-ESEEM spectra. On the left, chlorophyll *a* and phylloquinone structural composition of canonical Chl *a*-only binding PSI reaction centre (pdb id: 1JB0)^30^, representative of WL-adapted conditions, indicating the electron transport via the A-branch to the A_1A_ quinone acceptor (cyan dotted arrow) or via the B-branch to the A_1B_ quinone acceptor (red dotted arrow). The same for FR-PSI RC is presented in the central panel (pdb id: 9I9L)^26^, highlighting pigment substitution by Chl *f* (magenta). Phytyl chains are removed to improve clarity. On the right, model spin-echo decay and oop-ESEEM time traces originating from [P_700_^+^A_1A_^−^] (cyan lines) and [P_700_^+^A_1B_^−^] (red lines), based on previous studies on canonical PSI. Structure visualisation performed using PyMOL.

In order to shed light on the directionality of ET in FR-PSI, we here present the analysis of the photochemically generated chlorophyll-phylloquinone spin-correlated radical pairs [P_700_^+^A_1A/B_^−^], which serve as selective internal probes of ET pathways in PSI. This study combines Pulse and Time-Resolved (TR) Electron Paramagnetic Resonance (EPR) spectroscopy at 80 K on cells, thylakoid membranes and purified PSI complexes from C. *thermalis* PCC 7203 acclimated to both white light (WL) and FR illumination. The kinetic decay of the electron spin-echo amplitudes, the oscillation frequencies that dominate out-of-phase Electron Spin Echo Envelope Modulation (oop-ESEEM) spectra, together with complementary TR-EPR spectra recorded at Q-band, are diagnostic probes for the radical pairs populated on each PSI ET chain (Figure 1). Their combined analysis consistently supports the interpretation that bidirectional ET operates in FR-PSI, analogously to what observed in canonical Chl *a*-only systems, thereby providing a framework for further discussions concerning the energetics and redox properties of the RC cofactors in FR-PSI.

## 2. Materials and Methods

### 2.1 Sample preparation

*C. thermalis* PCC 7203 was grown in BG11 medium employing either WL (LED bulb, 3 W, 2700 K) or FR light (750 nm, FWHM 30 nm), as previously described.^16^ Thylakoid membranes (TM) purification and successive PSI isolation by sucrose gradient separation from WL- and FR- adapted cells were performed as previously described.^16,31^ The lower bands containing the PSI complexes of interest were pooled and concentrated by centrifugation-assisted dialysis (100 KDa cut-off). Sample purity was checked by UV-Vis spectroscopy and 6M Urea SDS-PAGE. All operations were conducted at 4 °C, in darkness or under dim green light. The obtained complexes at a concentration equivalent to 20-40 O.D. cm^−1^ at 680 nm (depending on the purification batch) were then frozen in liquid nitrogen and stored until use.

*C. thermalis* PCC 7203 cells, grown in WL or in FR light, were dark-adapted for 10 minutes, diluted with degassed glycerol to a final concentration of 60 % v/v and rapidly frozen in liquid nitrogen before EPR measurements. Dark-adapted WL- and FR-TM, as well as isolated PSI, were incubated on ice in the dark for 10 minutes following the addition of sodium ascorbate (20 mM). Degassed glycerol was then added (60 % v/v) after which the samples were transferred into EPR tubes and frozen in liquid nitrogen.

Photoaccumulation of the phylloquinone acceptors was obtained upon illumination of WL- and FR-PSI, preincubated with 20 mM sodium dithionite and 20 μM phenazine methosulfate (PMS) at pH 8, at 220 - 230 K either by broadband WL or by selective FR illumination centred at ∼780 nm, yielding negligible excitation below 725 nm. See **Figure S1** for a schematic representation of the photo-accumulation set-up and further information on the experimental conditions.

### 2.2 Spectroscopic measurements

Electron Paramagnetic Resonance (EPR) experiments were performed on a Bruker Elexsys E580 spectrophotometer operating at either X-band (9.7 GHz) or Q-band (34.0 GHz). For X-band pulse EPR measurements, the samples were loaded into quartz tubes (3 mm i.d., 4 mm o.d.) and inserted into a Bruker Flexline ER4118X-MD5 dielectric resonator operating in the overcoupled conditions. For Q-band Time Resolved-EPR (TR-EPR) experiments, samples were loaded into quartz tubes (2 mm i.d., 3 mm o.d.) and measured using a Bruker Flexline ER 5106 QT-W (TE_011_ mode) critically coupled resonator. Temperature was controlled by an ER 4118CFO nitrogen flow cryostat governed by an ITC Mercury unit (Oxford). Photoexcitation was provided by a Quantell Brilliant Nd:YAG pulsed laser equipped with a second harmonic module, generating 532 nm light pulses with a duration of approximately 10 ns.

X-band pulse EPR experiments were performed at T = 80 K, using a laser pulse energy of 2 mJ and a repetition rate of 50 Hz. Electron spin echo (ESE) was generated by a two-pulse microwave sequence: *h*ν-DAF-*p*_1_-τ-*p*_2_-τ-echo. The duration of the first and second (hard) pulses, *p*_1_ and *p*_2_, was adjusted to 12 ns and 24 ns respectively. ESE decay kinetics were recorded by fixing the inter-pulse delay (τ) at 140 ns while varying the delay after laser flash (DAF) from 0 to 60 μs (or from 0 to 35 μs for photoreduced samples). oop-ESEEM measurements were acquired by incrementing τ from 148 ns to 2000 ns by 8 ns steps while maintaining the DAF at either 0.3 μs or 20 μs. ESE intensities were obtained by integration of the time domain traces over a 100 ns time window covering the whole echo. During all experiments, the magnetic field was fixed at the maximum of the ESE signal as determined from a preliminary field-swept ESE measurement. A two-step phase cycling was applied to the first pulse *p*_1_. Small phase corrections on the real and imaginary quadrature channels were applied during data processing.

Q-band TR-EPR experiments were performed at T = 80 K, using a laser pulse energy of 5 mJ, a repetition rate of 10 Hz, a microwave power of 50 μW (26 dB attenuation, SuperQ FT bridge) and a magnetic field sweep of 10 mT with a step size of 0.03 mT. Direct EPR absorption/emission transients (without field modulation) were recorded in diode detection mode, digitized and summed by the internal SpecJet-III module of the EPR console after being amplified by an external DHPVA-200 voltage amplifier. Background signals were removed by subtracting traces recorded at magnetic field positions well outside the resonance region, thereby eliminating the intrinsic resonator response. The shown spectra were obtained by averaging the transient signals over the 0.6 - 0.8 μs time window following laser flash excitation. For direct comparison, spectra from different samples were referenced to a common microwave frequency of 34.0 GHz by applying the appropriate horizontal field shift.

## 3. Data Analysis

### 3.1 ESE decay analysis

ESE decay kinetic traces were fitted by a least-squares minimisation method using the Nelder-Mead simplex algorithm, employing a laboratory-written script in MATLAB (version R2024b), according to the following biexponential decay model:

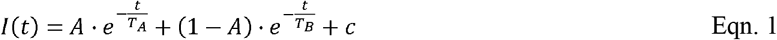

where *A* is the relative amplitude of the slow-decaying component, *T_A_* is the lifetime of the slow-decaying component, *T_B_* is the lifetime of the fast-decaying component, *c* is a constant accounting for small baseline offsets (constrained to remain below 2% of the initial signal) and *t* is the delay between the laser flash and the start of the microwave sequence (DAF). *T_A_* and *T_B_* were employed as global fitting parameters, treating datasets from WL and FR samples separately.

### 3.2 Out-of-Phase (oop)-ESEEM analysis

In spin-correlated radical pairs, under high-field approximation and considering ideal microwave (MW) pulses exciting the whole ensemble of spins, the oop-ESEEM response is dominated by electron-electron interactions, whereas contributions from nuclear hyperfine couplings are comparatively weaker and can be therefore neglected for simplicity. Following established theoretical treatments,^32–35^ the time domain oop-ESEEM can be approximated by:

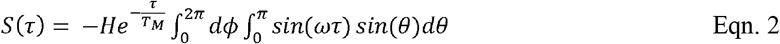

where *H* is the initial amplitude, *ω* is the ESEEM frequency, *τ* is the inter-pulse delay between the two pulses of the echo sequence and the exponential factor accounts for damping due to spin–spin relaxation processes, parameterized by the phase-memory time *T_M_*. Because the samples represent randomly oriented ensembles (powder samples), the observed oop-ESEEM signal corresponds to an average over all molecular orientations with respect to the external magnetic field, given by the double integral. The modulation frequency *ω* is directly correlated to electron-electron interactions among the spin correlated radical pair described by the expression 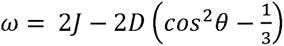, where *J* describes the isotropic exchange interaction, *D* describes the axially symmetric dipolar coupling and *θ* represents the angle formed between the direction of the external magnetic field and the direction of the principal axis of the dipolar interaction within the radical pair. While *J* typically shows exponential dependency on inter-spin distance, the dipolar coupling follows the point-dipole approximation and scales with the inverse cube of the separation distance according to:

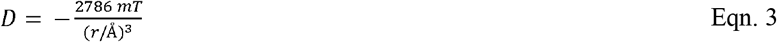

Where *r* denotes the effective distance between the centres of the spin density distributions of the two radical species, an approximation valid for sufficiently separated spins within the point dipole–point dipole regime. oop-ESEEM time domain traces were fitted to this model via a least-squares minimisation method based on the Nelder-Mead simplex algorithm. A parabolic offset was additionally introduced in the fitting model, as commonly seen in similar works. Fourier-Transformed spectra (imaginary component) were computed from both experimental and simulated time-domain traces. Prior to transformation, zero-filling, analytical reconstruction of the instrumental dead-time and subtraction of the parabolic offset were performed to increase quality. Uncertainties in the fitted parameters were estimated through the Cramér-Rao lower bounds theorem, as established in literature.^36^

All calculations were performed via a laboratory-written script operating in MATLAB (version R2024b).

### 3.3 TR-EPR spectra simulations

In contrast to oop-ESEEM spectra, which are mainly affected by dipolar interaction within the radical pair, and thus by the distance between the two spins, TR-EPR spectra are strongly influenced by the relative orientation of the principal axes of the ***g***-tensor frames associated with the chlorophyll and the quinone radicals, as well as by the principal axes of the dipolar interaction tensor.^37–39^ For a two-spin system (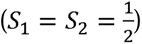), in the laboratory frame and neglecting the contributions of hyperfine couplings, the system Hamiltonian 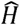 can be described as:

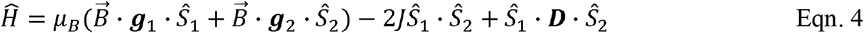

where *μ_B_* is the Bohr magneton, 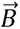 is the external magnetic field, *g*_1_ and *g*_2_ are the g-tensors of each spin, *J* is the isotropic exchange interaction and *D* the dipolar coupling tensor.

Q-band TR-EPR spectra were simulated by defining the orientations of the principal axes reference frames of all the involved tensors, as described in **Supporting Information**. For simplicity, hyperfine interactions were not explicitly included. TR-EPR simulations were performed using a laboratory-written MATLAB (version R2024b) script, taking advantage of the EasySpin toolbox.^40,41^

## 4. Results

### 4.1 Kinetics of ESE relaxation as a function of laser excitation delay

Photochemical charge separation in PSI rapidly (< 50 ps) populates the spin-correlated [P_700_^+^A_1_^−^] radical pair (RP), where P_700_^+^ is the terminal donor cation and A_1_^−^ the phyllosemiquinone acceptor anion.^42,43^ This radical pair (RP) gives rise to the intense and characteristic out-of-phase ESE signal that can be analysed by light-induced pulse EPR through its decay as a function of the delay after laser flash (DAF). The observed decay is primarily governed by spin relaxation processes, although it is also influenced by charge recombination reaction kinetics. At room temperature, [P_700_^+^A_1_^−^] undergoes rapid forward ET to iron-sulphur (Fe-S) centres on a nanosecond timescale, which substantially precludes detection by pulse EPR. In contrast, at T < 180 K, approximately half of the PSI reaction centres undergo charge recombination on the microsecond timescale rather than proceeding to downstream Fe-S reduction (at least in canonical PSI),^42,44^ which makes the process accessible to EPR investigation. Monitoring the [P_700_^+^A_1_^−^] decay kinetics at cryogenic temperature provides a convenient mean to observe the RP population on the two active ET branches of PSI, as they are typically characterised by neatly distinct decay lifetimes.^45–48^ **Figure 2** shows the comparison of the ESE decay kinetics measured at 80 K in dark adapted cells, thylakoid membranes and isolated PSI of the cyanobacterium *C. thermalis* PCC 7203 grown either in WL or FR-light. It can be straightforwardly appreciated that all samples in which the FaRLiP-associated reorganization has been elicited exhibit a faster decay compared to the corresponding WL-adapted counterparts, indicating a modified balance between the slow- and fast-relaxing RP populations.

**Figure 2.**
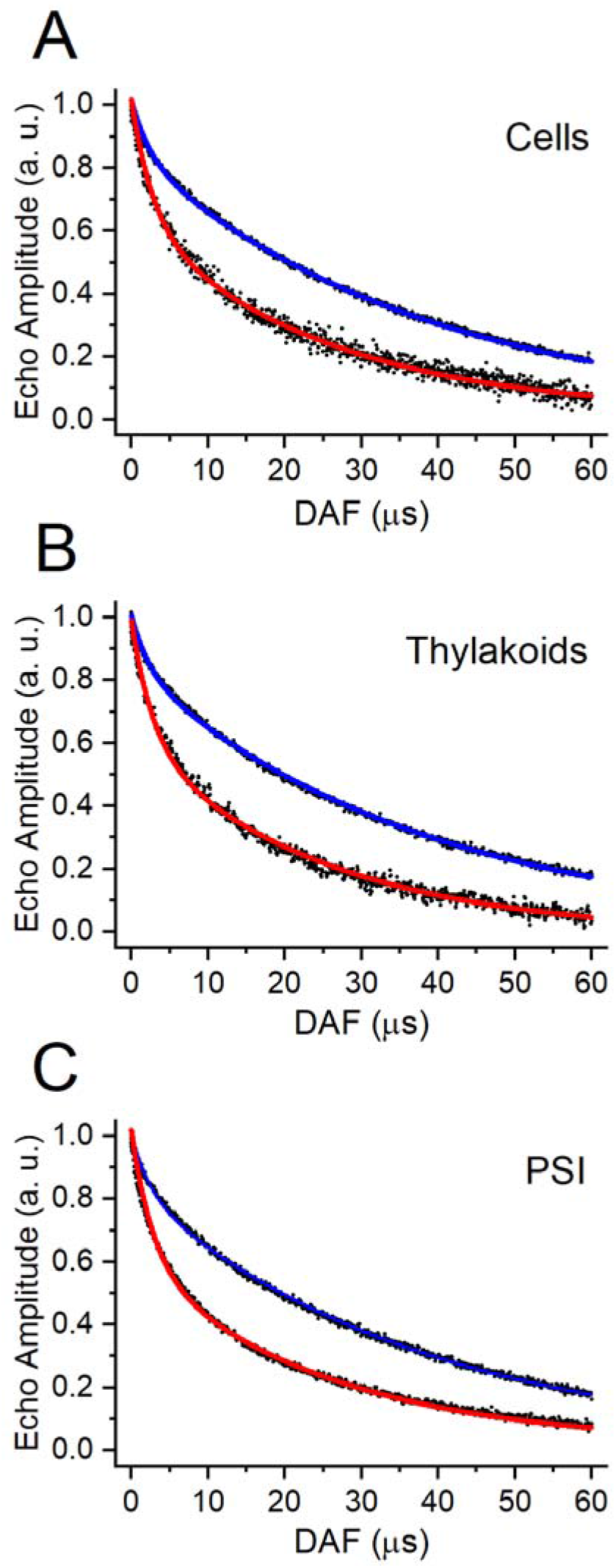
ESE decay kinetic traces as a function of the delay after laser flash (DAF) for dark-adapted **A:** cells; **B:** thylakoid membranes; **C:** isolated PSI. Experimental points are reported in black. Datasets were fitted via a biexponential decay model for WL samples (blue lines) and FR samples (red lines). Fitting parameters are reported in **Table 1**.

**Table 1.**
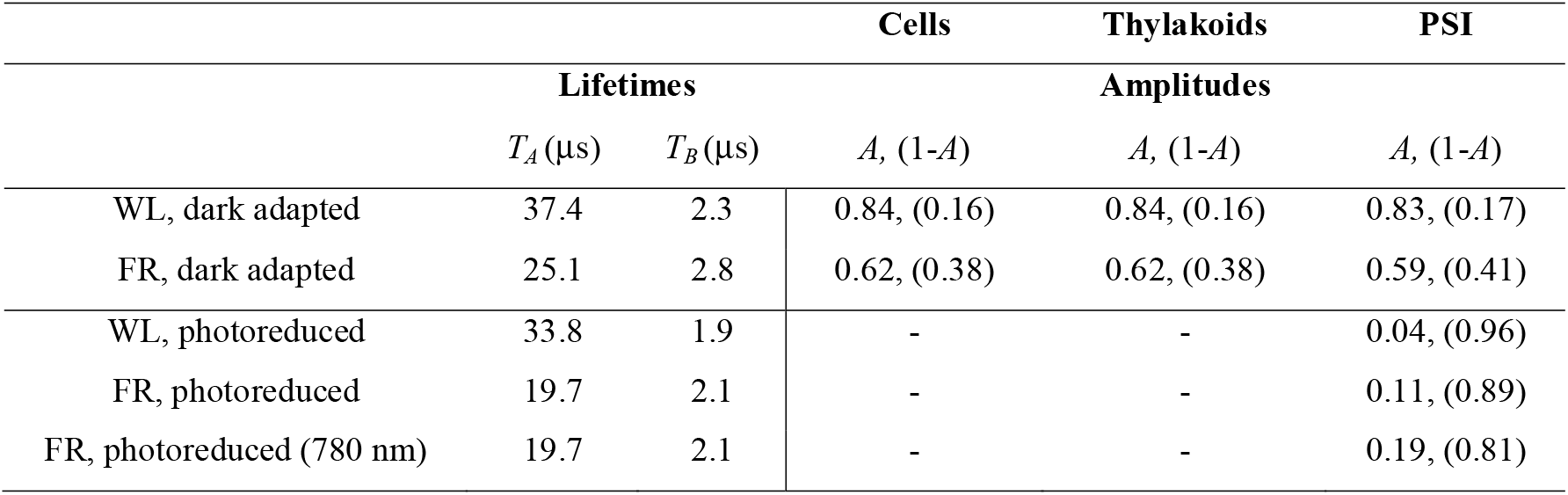
Fitting parameters of the ESE decay kinetic traces of dark-adapted and photoreduced samples. Fitting model: 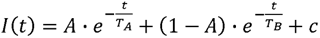.

To quantify the relative contributions of slow and fast decaying components, ESE decay traces were globally fitted via a biexponential decay model (see Section 3.1 for details). The resulting fitting parameters are reported in **Table 1**. In WL samples, the decay kinetics are dominated by a long-lived component with a lifetime *T_A_* = 37.4 μs, which accounts for 83-84 % of the total spin-echo amplitude. A minor contribution is given by a fast-decaying term (*T_B_* = 2.3 μs) contributing 16-17 % of the total amplitude. Comparable amplitude distributions are observed from intact cells to isolated PSI complexes, indicating that the relative efficiency of the underlying electron-transfer pathways is preserved upon membrane and complex isolation, thus excluding purification artefacts. These results are in good agreement with literature data obtained in dark adapted samples from non-FaRLiP photosynthetic organisms including *Synechocystis* PCC 6803 and *Chlamydomonas reinhardtii.*^46,48^

In contrast, FR-acclimated samples exhibit a markedly biexponential behaviour showing around 0.6 : 0.4 partition between the slow and the fast-decaying components. This trend is again consistently conserved across all FR-adapted samples. The lifetime of the slow decaying term in WL complexes is longer (37.4 μs) than in FR samples (25.1 μs). Although this difference might be considered significant, variations of several microseconds in the specific relaxation lifetimes have been reported for different non-FaRLiP organisms (spinach, green algae, cyanobacteria)^47^ as well as for the Chl-*d* containing cyanobacterium *A. marina*^49^, suggesting an influence of the specific protein environment on the relaxation processes. The most notable difference between WL- and FR-acclimated samples, however, lies in the relative amplitude of the fast-decaying component. In FR complexes, this contribution increases to ∼ 40% of the initial ESE amplitude, more than twice that observed under WL conditions. Given that the slow-decaying component has previously been associated with the [P_700_^+^A_1A_^−^] radical pair formed via the A-branch ET, whereas the fast-decaying component has been attributed to [P_700_^+^A_1B_^−^] (B-branch ET radical pair),^46–48^ these results indicate that both ET branches are photochemically active even after FaRLiP reorganisation.

**Figure 3A, B** shows the ESE relaxation of WL- and FR-adapted PSI which were subjected to prolonged broadband visible-light illumination under reducing conditions at 230 K, a treatment that, at least in canonical PSI, promotes the reduction of the Fe-S centres and the accumulation of singly-reduced A_1A_.^50,51^ For FR-PSI, an additional photoreduction procedure was applied using far-red excitation through a 780 nm long-pass filter that selectively excites the Chl *f* molecules in this system (**Figure 3C**). The efficacy of the photoreduction treatments was verified by monitoring the production of stable radical species, *i.e.* A_1_^−^ anion and reduced Fe-S centres, which give typical X-band EPR signatures (spectra presented in **Figure S2** and **S3**). Despite differences in illumination conditions and pre-irradiation times, all samples converge to a relaxation behaviour dominated by a fast component with a lifetime in the range of 1.9–2.1 μs, as summarised in **Table 1**. These findings agree with previous reports on canonical Chl *a*–binding PSI systems, where selective photoaccumulation of the A_1A_ radical suppresses the long-lived (*T_A_* = 25–40 μs) decay component, enabling preferential observation of the remaining radical-pair contribution associated with the B-branch ET pathway.^47,52^ Following photoreduction treatments, all samples exhibit systematically shortened decay lifetimes relative to dark-adapted conditions (**Table 1**). This can be explained considering the influence of reduced F_X_, since the presence of an additional unpaired electron in the proximity of A_1_ acceptor likely accelerates spin-relaxation dynamics.

**Figure 3.**
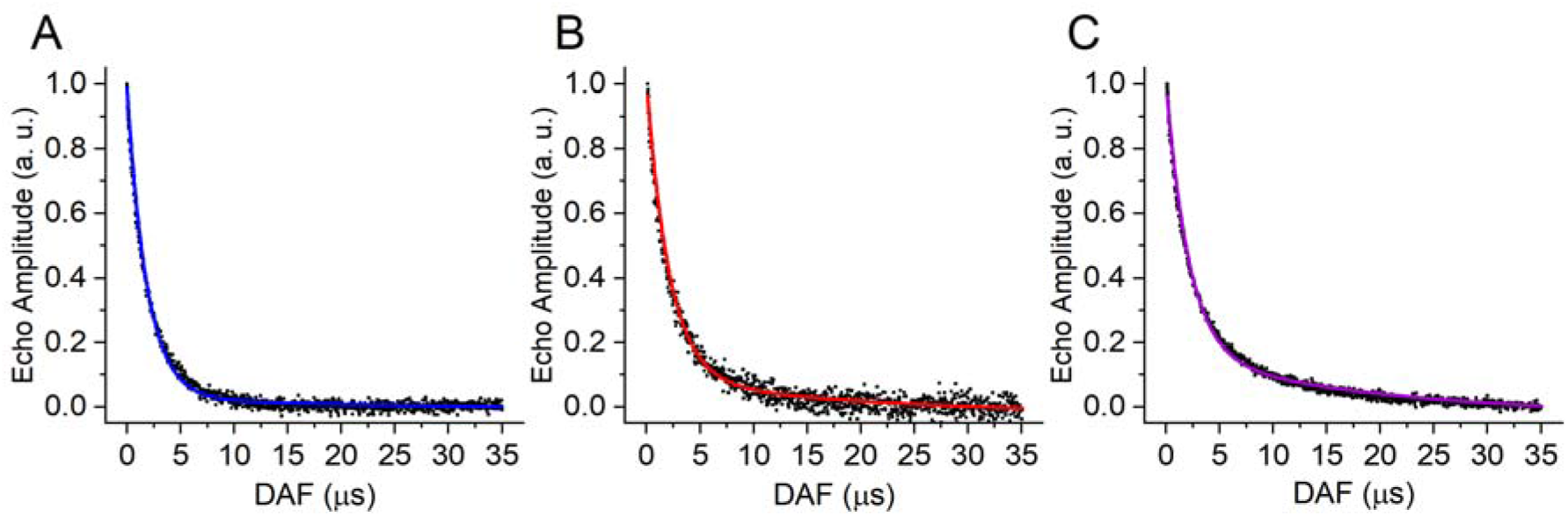
ESE decay kinetic traces of reduced and photoaccumulated samples from **A:** WL-PSI (data fitting marked with a blue line); **B:** FR-PSI under broad visible light (data fitting marked with a red line); **C:** FR-PSI under 780 nm filtered light (data fitting marked with a purple line). Fitting parameters are reported in **Table 1**.

### 4.2 Frequency modulation of the out-of-phase Electron Spin Echo Envelope Modulation (oop-ESEEM)

The above interpretation is further supported by the analysis of the oop-ESEEM frequency components, providing valuable information concerning the dipolar interaction between RP partners, from which the distance between the centroids of the unpaired electron spin density distributions associated to the chlorophyll cation and the quinone anion can be accurately estimated.^36,53^ Taking into account the well-established asymmetric spin distribution of the radical cation P_700_^+^ on the P_B_ half of the chlorophyll dimer,^54–56^ the P_700_^+^ - A_1A_^−^ distance can be predicted to be around 25.8 Å, larger compared to the 24.6 Å distance between P_700_^+^ and A_1B_^−^ (values obtained via analysis of PSI crystallographic structure, pdb id: 1JB0)^30^. These small but systematic differences in inter-spin distance translate into measurable variations in the dipolar coupling strengths and, consequently, in the oop-ESEEM modulation frequency. When recorded with sufficient signal-to-noise ratios, oop-ESEEM spectra are sensitive enough to resolve these differences, enabling discrimination between a lower modulation frequency (arising from [P_700_^+^A_1A_^−^]) and a higher one associated with [P_700_^+^A_1B_^−^] radical pair.

**Figure 4A** shows the oop-ESEEM spectra recorded from dark adapted WL- and FR-PSI, at two different laser–microwave delays after flash (DAF = 0.3 μs and 20 μs). The longer delay was chosen to ensure that the fast-relaxing contribution observed in the ESE decay kinetics, assigned to [P_700_^+^A_1B_^−^], is largely attenuated. In WL-PSI the oop-ESEEM frequency modulation is essentially unaffected by the DAF. In contrast, FR-PSI exhibits a faster modulation at DAF = 0.3 μs compared to DAF = 20 μs, the latter being also identical to oop-ESEEM traces for WL-PSI. This behaviour is consistent with a significantly larger contribution of a high-frequency modulation component in FR-PSI at short DAF, reflecting the enhanced population of the [P_700_^+^A_1B_^−^] radical pair, in agreement with its increased amplitude contribution observed in the biexponential ESE decay (**Table 1**). By contrast, WL-PSI, which displays a nearly monoexponential ESE relaxation behaviour, is dominated by an oop-ESEEM frequency component that remains substantially invariant with DAF, accounting for prevalent observation of [P_700_^+^A_1A_^−^]. **Figure 4B** compares oop-ESEEM traces at DAF = 0.3 μs for dark-adapted and photoreduced PSI samples from both WL- and FR-acclimated organisms. Regardless of growth conditions, oop-ESEEM spectra of samples subjected to photoreduction are characterised by a faster modulation frequency compared to dark adapted complexes, as expected because of the stronger dipolar couplings within the residual [P_700_^+^A_1B_^−^] radical pair.

**Figure 4.**
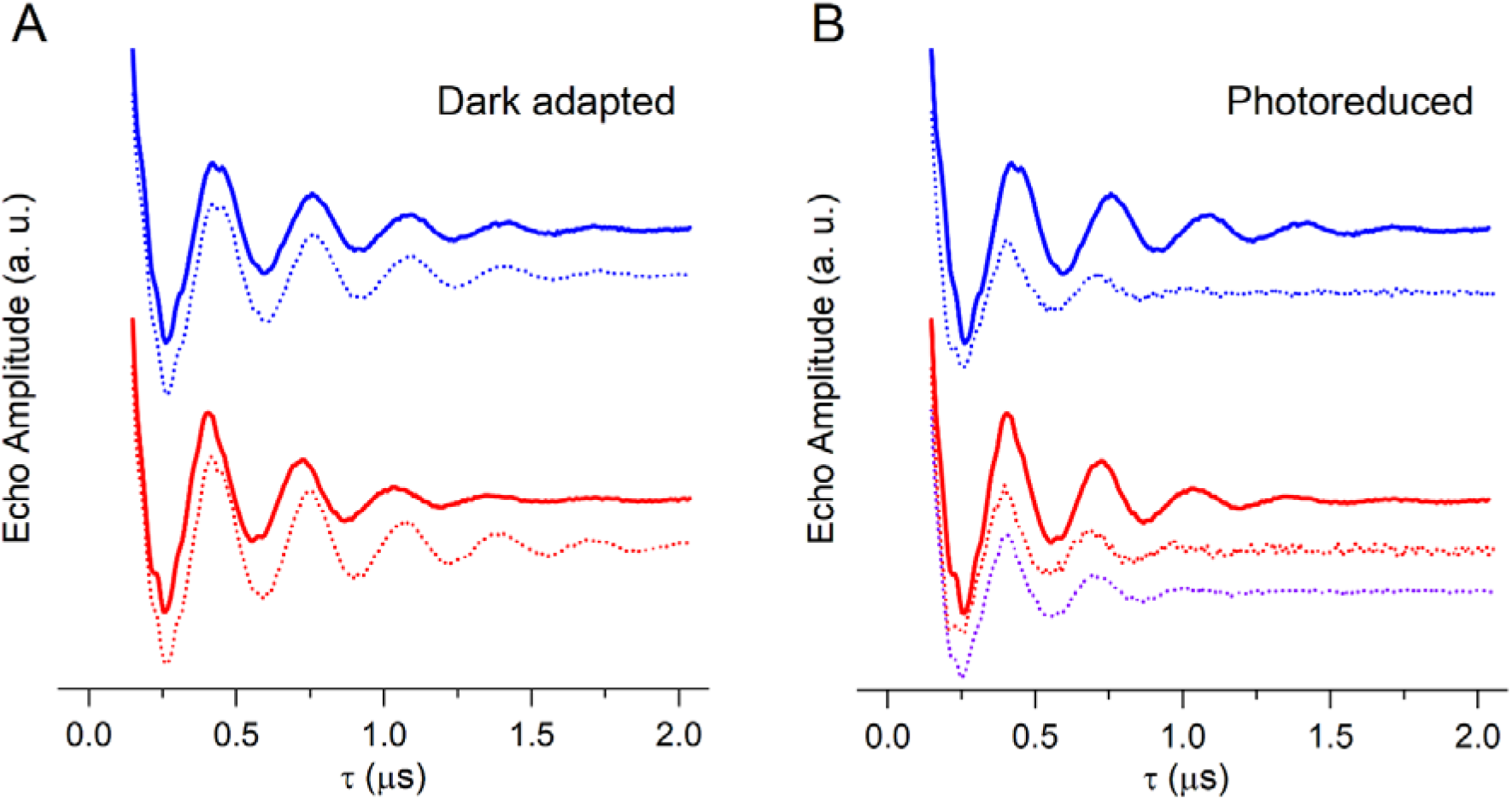
A: oop-ESEEM spectra of dark-adapted WL-PSI complexes (blue traces) and FR-PSI complexes (red traces) recorded at DAF = 0.3 μs (solid lines) or 20 μs (dotted lines). **B:** oop-ESEEM spectra recorded at DAF = 0.3 μs for reduced and photoaccumulated WL-PSI (blue dotted line), FR-PSI under visible light (red dotted line) and FR-PSI under 780 nm FR illumination (purple dotted line). For direct comparison, the spectra for dark adapted samples (solid lines) are also reported.

In support of these assignments, oop-ESEEM traces (in time domain and frequency domain) for WL- and FR-PSI dark-adapted samples were fitted following the procedure described in Section 3.2 (**Figure 5**). WL-PSI spectra recorded at DAF = 0.3 μs (**Figure 5A**) and at DAF = 20 μs (**Figure 5B**), satisfactorily described by a single-frequency model, yielded nearly identical values of the exchange and dipolar coupling parameters, *J* and *D* (**Table 2**). These parameters are also consistent with those obtained for FR-PSI at DAF = 20 μs (**Figure 5D**), indicating the observation of the same RP species under these detection conditions. In contrast, attempts to fit the FR-PSI spectrum at DAF = 0.3 μs using a single oscillation frequency were unsuccessful (**Figure S4**). A reliable description of the data required a two-frequency model with comparable amplitudes, one of which was constrained to the parameter set extracted from the long-DAF analysis at DAF = 20 μs (**Figure 5C**). This outcome confirms the coexistence of two distinct radical-pair contributions at short DAF in FR-PSI. In WL-PSI at DAF = 0.3 μs, fitting of the oop-ESEEM trace with two frequency components (**Figure S5**) gives results that are substantially indistinguishable from the data fitting with just a single frequency contribution. This remarks the fact that the fast-relaxing term in WL-PSI ESE kinetics, showing a weak amplitude, cannot be easily detected by oop-ESEEM, being masked by the larger signal associated to the slow-relaxing term. The analysis of the fitting parameters for WL-PSI oop-ESEEM yields coupling constants *J* = 0.5 μT and *D* = -168.6 μT, corresponding to an inter-spin distance of 25.44 Å, in excellent agreement with the structural models and consistent with dominant observation of charge separation along the A-branch of PSI.^35^ Very similar values (*J* = 0.5 μT; *D* = –170.9 μT; *R* = 25.33 Å) are obtained for FR-PSI at DAF = 20 μs, further supporting a common A-branch-dominated contribution for long DAF detection conditions. For the short-DAF FR-PSI data, the second (high) frequency component is characterized by significantly stronger coupling parameters, *J* = 3.0 μT and *D* = -196.1 μT, indicative of a reduced effective inter-spin distance (*R* = 24.19 Å) and thus consistent with a distinct radical-pair contribution associated with the B-branch ET pathway.^46–48^

**Figure 5.**
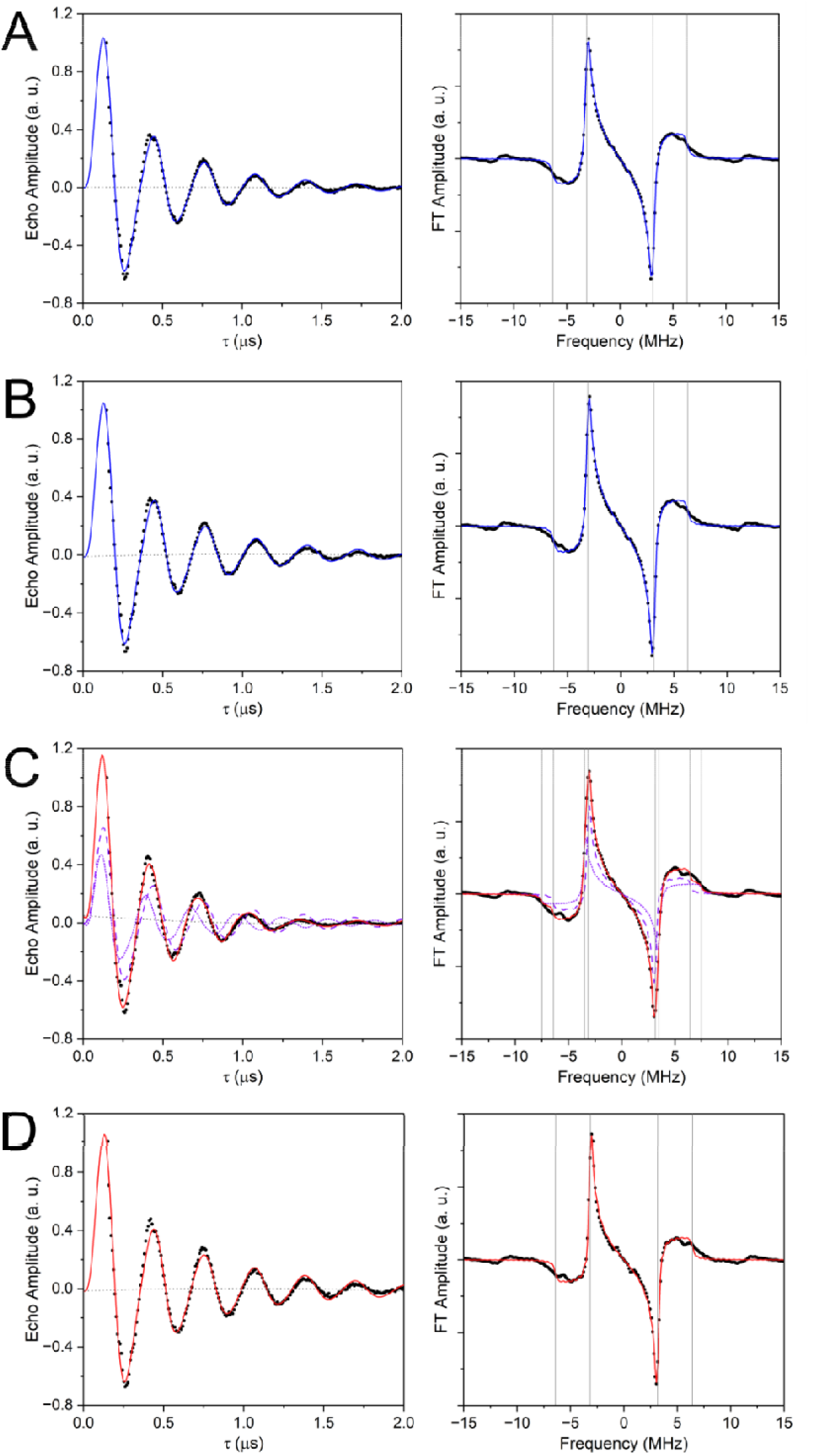
Fitting of oop-ESEEM spectra in time domain (left panels) and frequency domain (right panels) for dark adapted samples. **A**: WL-PSI at DAF = 0.3 μs; **B**: WL-PSI at DAF = 20 μs; **C**: FR-PSI at DAF = 0.3 μs; **D**: FR-PSI at DAF = 20 μs. Experimental points are reported with black dots, fitting traces are reported in coloured lines (blue for WL samples, red for FR samples). In panels **C**, the two frequency components are reported in purple dotted (high frequency) and dashed (low frequency) lines. On the left panels, parabolic baselines are shown with a grey dotted line. On the right panels, turning points at 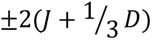 and 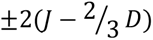 positions are marked by vertical lines.

**Table 2.** Fitting parameters of the oop-ESEEM traces of dark adapted and photoaccumulated PSI samples.

| | $T_M (\mu\text{s})$ | $J (\mu\text{T})$ | $D (\mu\text{T})$ | $R (\text{\AA})$ |
| --- | --- | --- | --- | --- |
| WL, DAF = $0.3\ \mu\text{s}$ | $0.71 \pm 0.02$ | $0.5 \pm 0.2$ | $-168.6 \pm 0.6$ | $25.44 \pm 0.03$ |
| WL, DAF = $20\ \mu\text{s}$ | $0.79 \pm 0.02$ | $0.5 \pm 0.2$ | $-167.8 \pm 0.6$ | $25.48 \pm 0.03$ |
| FR, DAF = $0.3\ \mu\text{s}$ (fast component) | $0.94 \pm 0.03$ | $3.0 \pm 0.2$ | $-196.1 \pm 0.6$ | $24.19 \pm 0.02$ |
| FR, DAF = $20\ \mu\text{s}$ | $1.01 \pm 0.04$ | $0.5 \pm 0.3$ | $-170.9 \pm 0.7$ | $25.33 \pm 0.04$ |
| WL photoaccumulated | $0.29 \pm 0.01$ | $3.6 \pm 0.4$ | $-187 \pm 1$ | $24.59 \pm 0.05$ |
| FR photoaccumulated | $0.30 \pm 0.01$ | $3.6 \pm 0.4$ | $-193 \pm 1$ | $24.31 \pm 0.05$ |
| FR photoaccumulated 780 nm | $0.32 \pm 0.01$ | $3.4 \pm 0.3$ | $-187.9 \pm 0.9$ | $24.54 \pm 0.04$ |

This is further confirmed by the fittings of oop-ESEEM spectra recorded for WL- and FR-PSI subjected to pre-illumination under reducing conditions (**Figure 6**) yielding dipolar coupling parameters within the ranges *J* = 3.4-3.6 μT, *D* = -(187-193) μT. These values correspond to inter-radical distances of 24.31–24.59 Å, in close agreement with the fast-frequency component extracted from dark-adapted FR-PSI spectra recorded at DAF = 0.3 μs, assigned to the [P_700_^+^A_1B_^−^] radical pair, consistently with almost mono-exponentially fast-component dominated ESE decay (**Figure 3**, **Table1**). Importantly, photoaccumulation induced in FR-PSI under either broadband visible or far-red excitation leads to essentially indistinguishable results, indicating that the specific excitation conditions do not alter the selective photoaccumulation of the A_1A_ radical.

**Figure 6.**
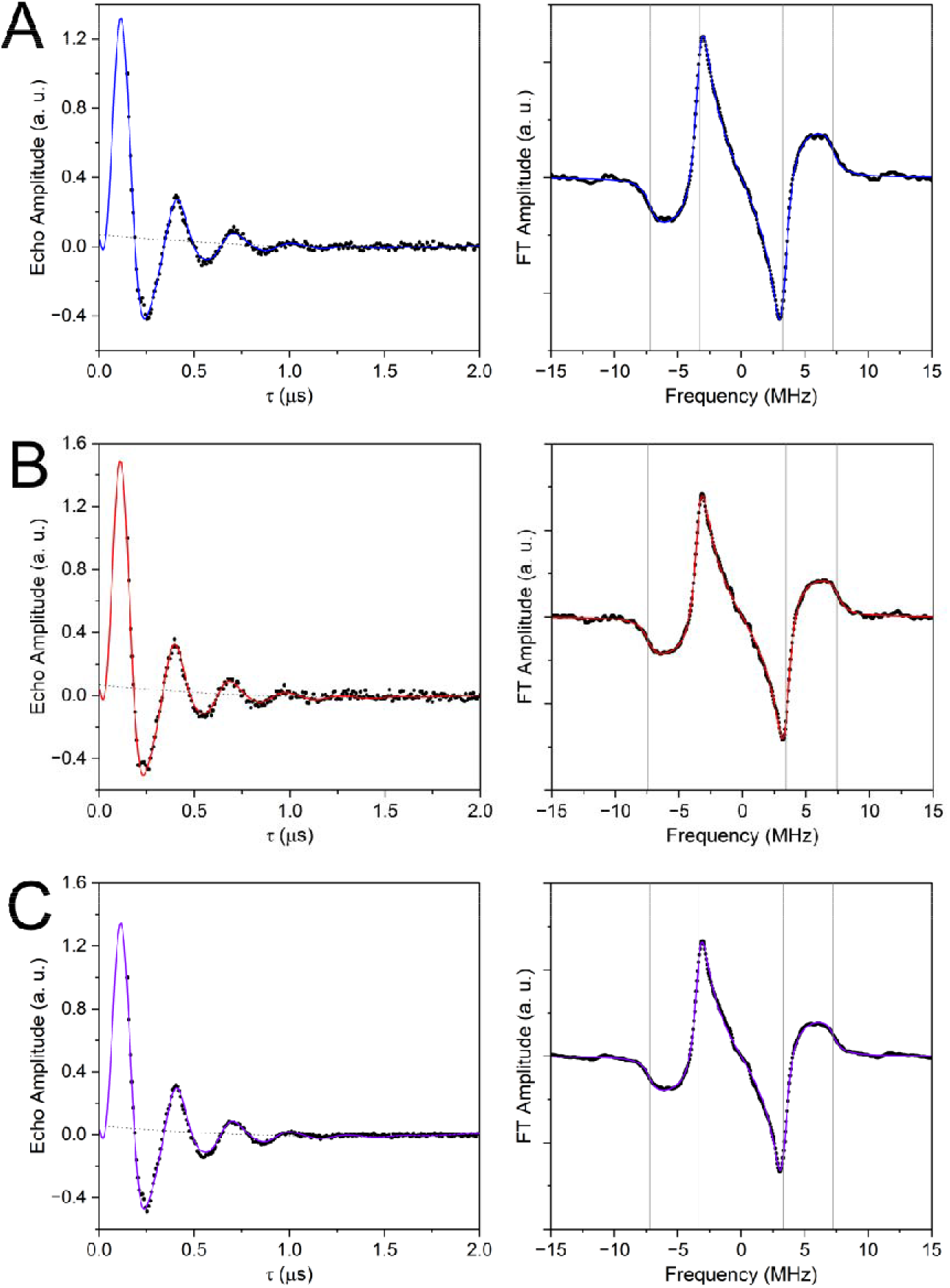
Fitting of oop-ESEEM spectra in time domain (left panels) and frequency domain (right panels) for reduced and photoaccumulated samples. **A**: WL-PSI; **B**: FR-PSI; **C**: FR-PSI reduced and photoaccumulated with 780 nm filtered light. Experimental points are reported with black dots, fitting traces are reported in coloured lines. On the left panels, parabolic baselines are shown with a grey dotted line. On the right panels, turning points at 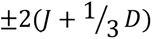 and 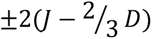 positions are marked by vertical lines.

### 4.3 Analysis of spin-polarised TR-EPR spectra

Whereas oop-ESEEM is highly affected by dipolar interactions, and thus by the relative distance between radical species, polarised TR-EPR spectra are considerably more sensitive to the relative orientation of RP partners, reflecting the mutual arrangement of the ***g***-tensor principal axes of P_700_^+^ and A_1A/B_^−^ as well as their orientation with respect to the principal axes of the dipolar tensor.^39,57^ In canonical PSI it was shown that the TR-EPR spectra associated with [P_700_^+^A_1A_^−^] and [P_700_^+^A_1B_^−^] radical pairs become readily distinguishable at sufficiently high microwaves frequencies,^38^ exhibiting nearly identical polarization patterns at low magnetic fields but an almost mirror-image polarization in the high-field region.^52,58^ Consequently, the polarization pattern of TR-EPR spectra provides a further sensitive probe of the electron-transfer branch responsible for charge separation in the PSI reaction centre.

**Figure 7** compares the Q-band TR-EPR spectra recorded for dark-adapted WL-PSI, dark-adapted FR-PSI, and reduced FR-PSI following photoaccumulation under broadband visible light illumination. Under all conditions, the low- and central-field regions of the spectra are remarkably similar, displaying an emissive polarization that evolves into an absorptive feature at the maximum of the positive TR-EPR signal. In contrast, the high-field region exhibits pronounced changes. In particular, the signal centred at approximately 1213 mT, together with its weaker high-field shoulder at 1213.5 mT, undergoes a progressive inversion of its polarization pattern, changing from emissive (absorptive) in WL-PSI to absorptive (emissive) in dark-adapted FR-PSI, with the inversion becoming even more pronounced after photoaccumulation. These changes in the high-field portion of the TR-EPR spectra are fully consistent with an increasing contribution of the B-branch charge-separated state. To better understand the origin of the spin polarisation inversion, TR-EPR spectra were simulated by defining the orientations of all the involved tensors, as well as the principal values of the ***g***-tensors and the spin-spin couplings (**Figure 8**, **Table 3**). Due to the similar structures and chemical environments of A_1A_ and A_1B_, the principal values of their ***g***-tensors were assumed to be identical and were taken from the literature.^52^ *J* and *D* coupling parameters were those determined from the fittings of the oop-ESEEM traces (**Table 2)**. The simulations therefore focused on the different relative orientations of the quinone and chlorophyll ***g***-tensors with respect to the dipolar interaction axis. Inspection of the relative orientations of the spin–spin dipolar axis with respect to the ***g***-tensor principal-axis systems of A_1A_ and A_1B_ reveals that the corresponding polar coordinates are highly similar, diverging less than 10°, between [P_700_^+^A_1A_^−^] and [P_700_^+^A_1B_^−^] (**Table 3**, **Figure 8**). In contrast, the description of the dipolar axis with respect to the chlorophyll ***g***-tensor frame shows large angle deviations between [P_700_^+^A_1A_^−^] and [P_700_^+^A_1B_^−^], especially for the polar angles *θ* that diverge by almost 50° (**Table 3**, **Figure 8)**. The preservation of the polarization pattern in the low-field region of the TR-EPR spectra, together with the pronounced polarization inversion at high magnetic fields, can therefore be rationalized by the nearly unchanged orientation of the dipolar tensor relative to the quinone ***g***-tensor and its markedly different orientation relative to the chlorophyll ***g***-tensor.

**Figure 7.**
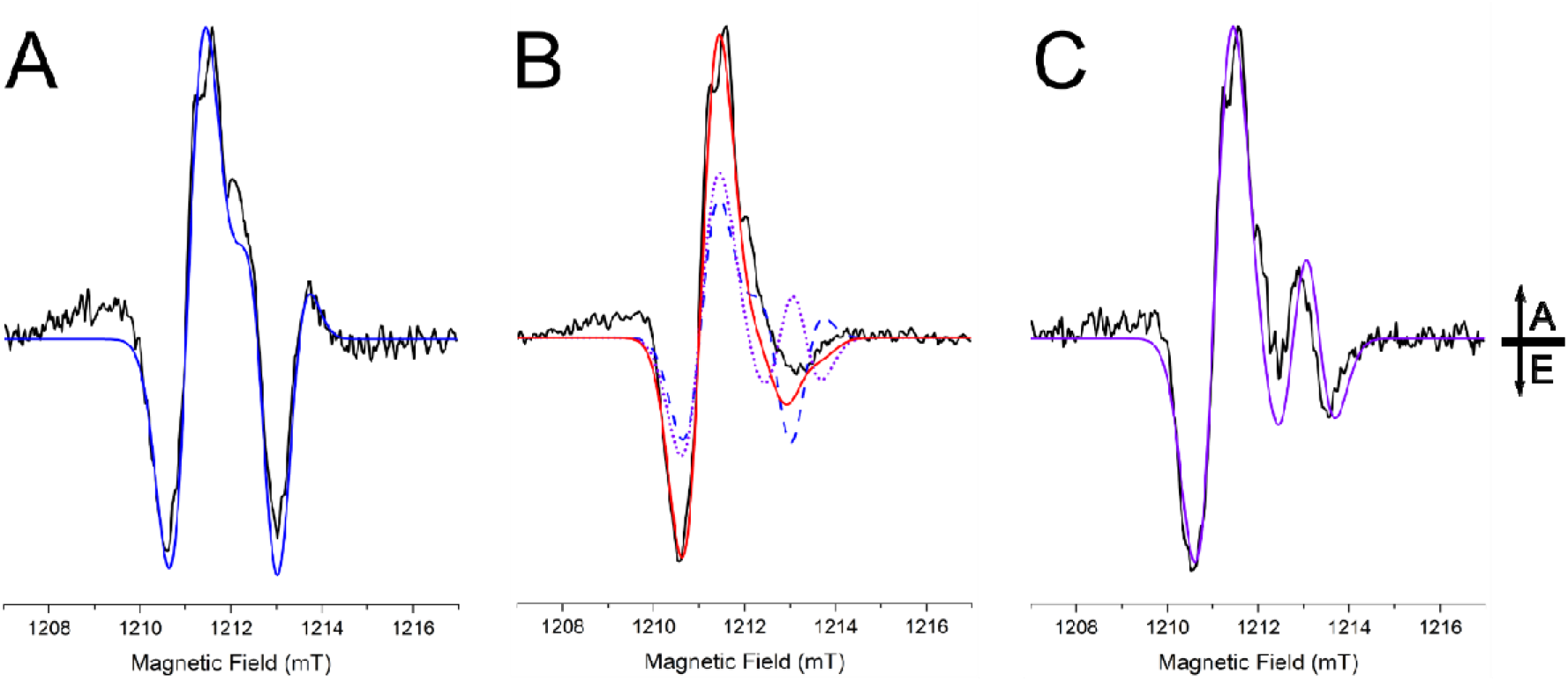
Q-band TR-EPR spectra of **A**: WL-PSI, together with its simulation (blue line) accounting for the prevalent radical pair [P_700_^+^A_1A_^−^]; **B**: FR-PSI, together with its simulation (red line) obtained as a sum of two contributions accounting for both [P_700_^+^A_1A_^−^] (dashed blue line) and [P_700_^+^A_1B_^−^] (dotted purple line); **C**: FR-PSI reduced and photoaccumulated under visible broadband light, together with its simulation (purple line) accounting for the prevalent radical pair [P_700_^+^A_1B_^−^].

**Figure 8.**
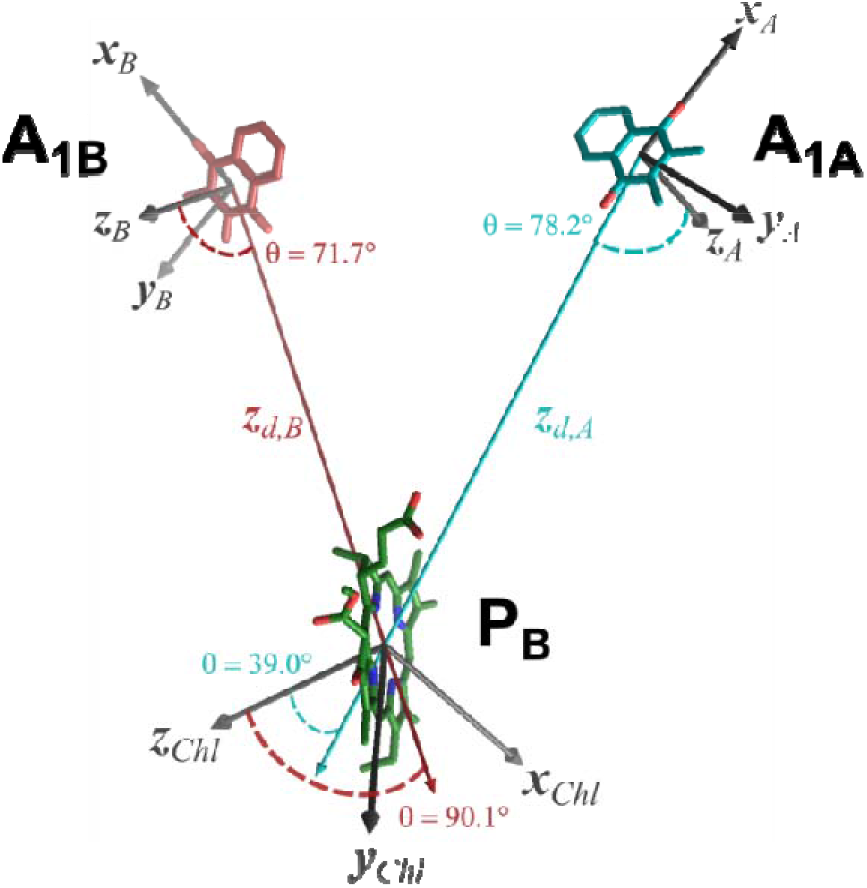
Schematic representation of all the tensor reference frames that are relevant for the simulations of TR-EPR spectra. (***x*_A_, *y*_A_, *z*_A_**) is the ***g***-tensor reference frame of A_1A_ quinone. (***x*_B_, *y*_B_, *z*_B_**) is the ***g***-tensor reference frame of A_1B_ quinone. (***x*_Chl_, *y*_Chl_, *z*_Chl_**) is the ***g***-tensor reference frame of chlorophyll in position P_B_. **z_d,A_** and **z_d,B_**represent the spin-spin dipolar axes for [P_700_^+^A_1A_^−^] and [P_700_^+^A_1B_^−^] respectively. Structural visualisation using PyMOL.

**Table 3.** Simulation parameters of the TR-EPR spectra, mainly retrieved from literature data^52^.

| | $g$-tensor principal values | | | Polar coordinates of dipolar axis with respect to $P_{700}^+$ | | |
| --- | --- | --- | --- | --- | --- | --- |
| | $g_x$ | $g_y$ | $g_z$ | | $\theta$ | $\phi$ |
| $P_{700}^+$ | 2.00322 | 2.00277 | 2.00246 | $[P_{700}^+ A_{1A}^-]$ | 39.0 | 62.0 |
| $A_1^-$ | 2.00636 | 2.00520 | 2.00230 | $[P_{700}^+ A_{1B}^-]$ | 90.1 | 54.8 |
| | Couplings and Linewidths | | | Polar coordinates of dipolar axis with respect to $A_1^-$ | | |
| | $J$ (μT) | $D$ (μT) | Linewidth (mT) | | $\theta$ | $\phi$ |
| $[P_{700}^+ A_{1A}^-]$ | 0.5 | -168 | 0.85 | $[P_{700}^+ A_{1A}^-]$ | 78.2 | 176.4 |
| $[P_{700}^+ A_{1B}^-]$ | 3.5 | -196 | 0.90 | $[P_{700}^+ A_{1B}^-]$ | 71.7 | 168.6 |

A satisfactory simulation of the TR-EPR spectrum of dark-adapted WL-PSI is obtained by considering, as approximation, only the formation of the radical pair [P_700_^+^A_1A_^−^], in agreement with the oop-ESEEM analysis (**Figure 5**). Considering a combination of two RPs, did not result in any significant improvement in the quality of the simulation (**Figure S6**). Conversely, the spectrum of dark-adapted FR-PSI is accurately reproduced only when weighted contributions from both the [P_700_^+^A_1A_^−^] and [P_700_^+^A_1B_^−^] radical pairs with almost equal amplitudes are considered, consistent with the two-component model required to fit the corresponding oop-ESEEM data recorded at DAF = 0.3 μs. Notably, the superposition of the two simulated radical-pair spectra results in substantial cancellation of spin polarization in the 1213–1214 mT region, reproducing the pronounced reduction in TR-EPR intensity observed experimentally for dark-adapted FR-PSI relative to WL-PSI. Finally, the TR-EPR spectrum of reduced and photoaccumulated FR-PSI was satisfactorily simulated considering only the presence of the B-branch radical pair [P_700_^+^A_1B_^−^], in complete agreement with the ESE decay kinetics and oop-ESEEM analysis.

## 5. Discussion

### 5.1 Bidirectional electron transfer in Far-Red light adapted PSI and implications for reaction centre organisation

The EPR characterization of the spin-correlated [P_700_^+^A_1_^−^] radical pair at cryogenic temperatures provides a direct experimental probe of the two photochemically active electron-transfer branches of PSI. In the present work, the population of the two charge-separated states were independently assessed using three complementary spectroscopic observables: (i) the ESE relaxation kinetics as a function of the delay after laser excitation, which discriminate the two radical pairs through their distinct relaxation lifetimes; (ii) the dipolar coupling extracted from oop-ESEEM measurements, which is a sensitive probe of the inter-spin distance between P_700_^+^ and A_1_^−^; and (iii) the polarization pattern of Q-band TR-EPR spectra, which is primarily determined by the relative orientations of the P_700_⁺ and A_1_^−^ ***g***-tensors with respect to the dipolar interaction tensor.

The remarkable agreement among these three experimental approaches demonstrates that in the dark-adapted FR-PSI, at low temperature, both the [P_700_^+^A_1A_^−^] and [P_700_^+^A_1B_^−^] radical pairs (RP) are observed with comparable amplitudes, providing compelling evidence that electron transfer proceeds through both cofactor branches in the FaRLiP-remodelled PSI reaction centre. The observation of spin-correlated EPR signals imposes stringent mechanistic constraints on the assignment of the electron-transfer pathways. Spin polarization is maintained only when the observed RP is rapidly and directly populated from the immediate ET precursor, thereby retaining the singlet spin correlation generated upon primary photochemical charge separation. Any RP formed through a slower secondary ET process would undergo substantial loss of spin correlation, leading to severe attenuation - or complete disappearance - of the characteristic polarized EPR response.^32,59^ As a consequence, it can be concluded that [P_700_^+^A_1A_^−^] and [P_700_^+^A_1B_^−^] observed in FR-PSI are directly formed from the immediately upstream pairs, [P_700_^+^A_0A_^−^] and [P_700_^+^A_0B_^−^]. Alternative mechanisms involving secondary population of one quinone from the opposite branch are difficult to reconcile with the present observations. In canonical PSI, population of A_1A_^−^ *via* A_1B_^−^ has been proposed either involving F_X_ as an intermediate^60^ or potentially by-passing it. However, the F_X_-mediated inter-quinone ET lifetime was predicted to be several nanoseconds,^60^ which is sufficiently slow to lead to a substantial suppression of the spin-correlated ESE signal. Furthermore, because of the very rapid magnetic relaxation within iron-sulphur clusters, the resulting EPR signal would not only be damped but also significantly distorted, resulting in a EPR spectrum dominated by the residual spin polarisation of P_700_^+^,^39,59^ rather than by the characteristic radical-pair pattern observed experimentally (**Figure 7**). Direct electron transfer between A_1B_^−^ and A_1A_ without involvement of F_X_ is similarly unlikely to account for the observed signals. Even under the most favourable Marcus conditions ΔG^0^ = -λ (the latter being the reorganisation energy),^61^ the maximum transfer rate predicted for the 15.2 Å edge-to-edge separation between the two quinones is in the order of 1 μs^−1^, substantially slower than the F_X_-mediated ET and therefore incompatible with preservation of the observed spin correlation. Thus, it can be concluded that the [P_700_^+^A_1A_^−^] and [P_700_^+^A_1B_^−^] radical pairs detected in FR-PSI originate from independent primary charge separation reactions occurring along the A- and B-branches respectively, which are both active in ET reactions, according to the bidirectional electron transfer mechanism. More broadly, these findings indicate that bidirectional ET is not a peculiarity of canonical Chl *a*-binding PSI, but rather a conserved functional feature maintained even after the extensive structural and pigment remodelling associated with FaRLiP.

Regarding the identity and arrangement of the RC pigments, the simplest interpretation of the present results is that the FR-PSI retains an approximately symmetric functional organization. The EPR experiments presented here do not directly identify whether the individual RC chlorophyll-binding sites are occupied by Chl *a* or Chl *f*. Nevertheless, they impose stringent constraints on any mechanistic model describing primary charge separation. Recent cryo-EM structures of FR-PSI have proposed an asymmetric pigment arrangement in which Chl *f* occupies exclusively the A_-1B_ (eC-A2) site.^26^ Within this model, primary charge separation is predicted to stem from the A_-1B_ cofactor, generating the initial [A_-1B_^+^A_0B_^−^] charge separated state, evolving into the final [P_700_^+^A_1B_^−^] radical pair. Such a mechanism requires Chl *f* at A_-1B_ site to function as an efficient electron donor. It has been proposed that this is enabled by the conserved FaRLiP-specific residues Cys463 and Phe459 (numbering according to *T. elongatus*), which generate a localized negative electrostatic potential near the C2 formyl group of Chl *f*, thereby shifting its redox potential toward more negative values.^26^ Moreover, the FaRLiP-specific residue motif Val/Ile666 – Ser667 – Thr673 in PsaB2 was found to be the cause of a ∼5° rotation of the Chl *a* in position A_0B_ in FR-PSI compared to WL-PSI,^62^ which was suggested to increase the electronic coupling between A_-1B_ and A_0B_ based on DFT calculations and further supporting an almost unidirectional ET along the B branch.^26^ While these structural observations provide a plausible framework for enhanced B-branch activity, they do not readily account for the spectroscopic evidence presented here.

It can be argued that since EPR experiments were performed at low temperature with unselective excitation at 532 nm, a significant fraction of ET through the A-branch may be promoted by direct excitation of Chl *a* cofactors in the RC, thereby biasing the observed ET directionality ratio in favour of this branch. This however would imply that charge separation stemming from the directly excited Chl *a* would kinetically outcompete energy equilibration within the RC itself and with the long wavelength light harvesting Chls too. Although at present there is no direct kinetic evidence in this regard at cryogenic temperatures, this appears to be an unrealistic scenario especially considering the insertion of Chl *f* at the A_-1B_ site in the RC, which should act as a deep energy sink within it. At the same time, ultrafast spectroscopic studies of FR-PSI at room temperature have demonstrated that, upon visible-light excitation, energy transfer from Chl *a* to Chl *f* consistently precedes energy transfer to the RC.^20^ Consequently, excitation with either visible or far-red light is expected to have little influence on the ET process, as charge separation occurs only after excitation has been funnelled from the antenna Chl *f* pigments to the RC, irrespective of the excitation wavelength. In addition, steady-state fluorescence measurements at cryogenic temperatures have shown that fluorescence predominantly originates from Chl *f* regardless of the excitation wavelength.^10^ These findings support efficient energetic coupling between Chl *a* and Chl *f*, with excitation being rapidly transferred to the Chl *f* antenna before delivery to the RC.

Collectively, these observations indicate that both ET branches participate in photochemistry in FaRLiP PSI. The formation of two distinct spin-correlated RP further implies that primary charge separation must originate from a common excited precursor, thereby pointing at the cofactors composing P_700_ (P_A_/P_B_) as those acting as primary donors. Within a purely excitonic description of the RC, however, the presence of a Chl *f* molecule at the A_-1B_ site introduces a lower-energy state localized on this pigment. Especially at cryogenic temperatures, such a state would act as an energetic trap for excitation within the RC, substantially reducing the population of the excited state of P_700_ (P_700_*) and, consequently, its ability to initiate primary photochemistry. For P_700_* to maintain a significant population, it is therefore necessary to invoke additional electronic interactions beyond the simple excitonic model. One attractive possibility is to consider mixing of P_700_ excitonic interactions with charge transfer (CT) states, giving rise to CT-mixed states that are at least iso-energetic with the electronic transitions of Chl *f*, as suggested by Tros *et al.*^63^ In this scenario, excitation energy reaches the CT-mixed states of P_700_*, from which primary photochemistry along the two ET branches can be subsequently triggered. It has to be noted that the involvement of low-laying CT mixed states in the chlorophylls coordinated inside the RC has been already proposed in order to explain the functionality of canonical PSI under far-red illumination (also at temperatures as low as 8 K),^64,65^ which is known to harbour long-wavelength antenna Chls *a* (red chlorophylls) that also represent traps competing for RC excitonic localisation.^66,67^ Provided that the structures of these low-energy CT-mixed states is conserved in FR-adapted PSI, this may allow P_700_ to become effectively the lowest energy state in the RC and hence the site of primary charge separation. This would in turn be compatible with bidirectional electron transfer even in the presence of asymmetric cofactor composition. Yet, under this hypothesis, which will require further experimental confirmation, the Chl *f* molecule at the A_-1B_ position would act as an electron acceptor, rather than the primary donor.

### 5.2 Tuning of the redox-active cofactor properties

Even in face of the substantial conservation of the bidirectional mechanism, FR-PSI shows some significant characteristics which are distinct from what observed in the corresponding WL-PSI, and, more generally, in canonical PSI. The most striking difference is the clear detection of the spin-polarised EPR signal arising from both [P_700_^+^A_1A_^−^] and [P_700_^+^A_1B_^−^] with almost equal amplitudes under ambient redox conditions in dark adapted FR-PSI, while in canonical PSI the observation of [P_700_^+^A_1A_^−^] typically dominates. Only upon F_X_ pre-reduction^46–48^, or its removal by biochemical treatments,^68^ the two radical pairs are observed in almost equal proportion in canonical PSI. This behaviour was explained in terms of asymmetric driving forces for A_1_^−^ oxidation by F_X_ on the two ET chains, arising from branch-specific differences in quinone redox potentials.^28,29^ It was proposed that, in most RCs, A_1A_^−^ oxidation by F_X_ is endergonic and therefore becomes inhibited at cryogenic temperatures, promoting the observation of its spin-correlated radical pair, followed by charge recombination. Conversely, A_1B_^−^ oxidation by F_X_ is mostly exergonic leading to fast population of the irreversible [P_700_^+^F_A/B_^−^] state at cryogenic temperature. Blocking electron transfer by chemical pre-reduction of F_X_ makes then the radical pair [P_700_^+^A_1B_^−^] observable under repetitive flash detection methods together with [P_700_^+^A_1A_^−^].^48^ After prolonged photochemical reduction leading to complete accumulation of A_1A_^−^, only the [P_700_^+^A_1B_^−^] radical pair is observed, instead.^52^

The enhanced [P_700_^+^A_1B_^−^] contribution in FR-PSI can arise from two non-mutually exclusive causes. First, a redistribution of primary charge-separation probability between the two ET branches, induced by Chl *f* in A_-1B_, could favour the PsaB-coordinated pathway. Moreover, as already discussed, specific alterations in the A_0B_ binding pocket cause a 5° tilt of the cofactor changing its interaction with the Chl at position A_-1B_, possibly incrementing the probability of B-branch utilisation as well. In this respect, there is extensive experimental evidence that altering A_0_ coordination strongly affects ET directionality.^46,69–71^

Second, the thermodynamic landscape of downstream ET reactions may be altered, so that oxidation of A_1B_^−^ by F_X_ becomes less exergonic - or even endergonic - thereby increasing the lifetime and detectability of the B-branch RP. Such a shift could result from changes in the redox potential of A_1B_, F_X_ or from a combination of both effects. Notably, recent high-field pulse EPR studies on the green alga *Chlorella vulgaris* have reported simultaneous detection of [P_700_^+^A_1A_^−^] and [P_700_^+^A_1B_^−^] at cryogenic temperature with comparable amplitudes, that was consistently interpreted in terms of an endergonic A_1B_^−^ oxidation caused by an increased A_1B_ redox potential.^72^ In the same work, the green alga *Scenedesmus obliquus* showed even more pronounced signals attributed to [P_700_^+^A_1B_^−^] compared to [P_700_^+^A_1A_^−^].^72^ Although these measurements were performed on almost fully deuterated samples, which may influence spin and ET dynamics, they nevertheless indicate that the coexistence of both radical pairs is not exclusive to *C. thermalis* FaRLiP-reorganised photosystems but may also occur, albeit less frequently, in canonical PSI. Inspection of PsaB2 sequences and available FR-PSI structural models reveals several conserved amino-acid substitutions in the vicinity of the A_1B_ phylloquinone, the most significant being the replacement of Ser672 with Thr in FR-PSI.^62^ It is worth noting that this residue is also considered to be part of the FaRLiP-specific A_0B_ binding motif. Previous analysis of site-directed mutants of the Ser672 residue (substituted by Cys) in canonical PSI resulted in a slowdown of the A_1B_^−^ oxidation rate by F_X_ in the B-branch.^70^ Hypothesizing that this decrease in ET rapidity is the result of a higher redox potential of A_1B_ caused by Ser672 replacement, a similar situation can be qualitatively proposed even for the FaRLiP specific Ser to Thr exchange. At the same time, Fourier-Transform infrared difference spectroscopy across multiple cyanobacterial lineages, including *C. thermalis* PCC 7203 grown under both WL and FR conditions, has revealed a high degree of conservation in vibrational features associated with A_1_ binding pockets, particularly those assigned to quinone carbonyl stretching modes.^73^ However, the experimental conditions employed in the study appeared to favour the monitoring of the A_1A_ quinone. Therefore, small difference associated with A_1B_ environment might have escaped detection.

Finally, since F_X_ is coordinated at the interface between the PsaA and PsaB subunits, both of which are replaced by FaRLiP-specific paralogs, the associated backbone rearrangements and side-chain substitutions may alter the local electrostatic environment, thereby modulating the redox properties of the Fe-S cluster. A recent computational study based on cryo-EM structures of FR-PSI from the cyanobacteria *Halomicronema hongdechloris* and *Fischerella thermalis* PCC7521 acclimated to FR- light highlighted changes in the redox potentials of the Fe-S clusters.^74^ In particular, the midpoint redox potential of F_X_ was consistently predicted to be lower (*i.e.* more negative) in FR-PSI than in the corresponding WL-PSI complexes. This would in turn cause forward ET from A_1_ to F_X_ to become thermally activated on both branches in FR-PSI, thereby providing a plausible explanation for the observation of both spin-correlated radical pairs at cryogenic temperatures. Although this hypothesis is consistent with the present TR-EPR observations, direct experimental verification of the redox properties of F_X_ and of the temperature dependence of electron transfer in FR-PSI will be required to establish this mechanistic picture.

Collectively, these considerations can be taken as an indication that the tuning of A_1B_ redox properties, and perhaps those of F_X_ too, might be required to maintain a sufficient driving force for phylloquinone reduction and/or to avoid back-reactions leading to the repopulation of upstream cofactors at ambient temperatures. This would represent an intriguing fine-tuning of the FR-adapted PSI ET chains extending beyond the RC and the low-energy antenna states. However, definitive quantification of these energetic reconfigurations will require further targeted spectroscopic and computational investigations aimed at resolving the subtle energetic landscape modifications of FR-PSI RC.

## 6. Conclusions

Extensive characterisation of electron transfer within the FR-PSI reaction centre using complementary EPR methodologies provides a consistent description of the photogenerated [P_700_^+^A_1_^−^] spin-correlated radical pair. The combined analysis of ESE decay kinetics, oop-ESEEM modulation frequencies and Q-band TR-EPR polarization patterns support the occurrence of bidirectional electron-transfer according to which both the A- and B-branches of FR-PSI are photochemically competent.

The concomitant detection of the [P_700_^+^A_1A_^−^] and [P_700_^+^A_1B_^−^] radical pairs requires that primary charge separation originates at the level of P_700_, followed by ultrafast branching toward both electron-transfer chains. In this framework, the proposed structural asymmetry of FR-PSI associated with the incorporation of a single Chl *f* at the A_-1B_ position does not disrupt the intrinsic capability of the reaction centre to undergo photochemical charge separation on two parallel cofactors pathways. Importantly, the present results do not support a scenario in which Chl *f* acts as a unique primary electron donor. Instead, primary photochemistry would be required to initiate from P_700_, with Chl *f* most plausibly acting as a spectral and energetic modulator within the antenna–RC interface rather than as a branch-selective trigger of charge separation. Compared to WL-PSI, the increased relative contribution of the [P_700_^+^A_1B_^−^] radical pair in FR-PSI can be rationalized in terms of a modified energetic landscape on the acceptor side, consistent with a modest increase in the effective redox potential of A_1B_ and/or subtle tuning of the downstream driving forces toward F_X_. Collectively, changes in the nature of the ET cofactors and their interactions with the protein scaffold could shift the balance between the two electron-transfer branches utilisation without altering the fundamental architecture of bidirectional charge separation in PSI.

## Supporting Information

Description of the photoreduction procedure and setup; details on TR-EPR spectra simulations; CW-EPR spectra on quinone radicals and reduced Fe/S centres; additional oop-ESEEM fittings; additional TR-EPR spectra simulations.

## Acknowledgments

APC, SS, and DC acknowledge funding from the EU through MUR, for the project “Extending the red limit of oxygenic photosynthesis: basic principles and implications for future applications”, PRIN20224HJWMH (CUP B53D23015880006/C53D23004620006). AC acknowledges support from the Chemical Complexity C2 project (University of Padova, Dept. of Chemical Sciences). AA acknowledges financial support by the University of Padova (P-DiSC 2025).

## Notes

The authors declare no conflicts of interest of any kind.

## SUPPORTING INFORMATION

### Additional Material and Methods

#### Reduction and Photoaccumulation Treatments

Reduction and photoaccumulation treatments, allowing for the reduction of iron-sulphur centres and quinones (mainly one-electron reduction of A_1A_)^1,2^ were performed on isolated WL- and FR-PSI complexes as follows. Samples were incubated on ice in the dark for 5 minutes after the addition of sodium dithionite and phenazine methosulfate (PMS) at pH 8. Degassed glycerol was subsequently added (60 % v/v), resulting in final concentrations of 20 mM sodium dithionite and 20 μM PMS. Under these conditions, F_A_ and F_B_ are expected to be reduced by the action of dithionite in the dark. Further reduction of F_X_ and A_1A_ was achieved by low temperature photoaccumulation. Samples were immersed in a glass dewar containing an EtOH/CO_2_ cooling bath at 220 - 230 K and illuminated. WL-PSI and FR-PSI were photoreduced with broad-spectrum visible light through a combination of a 250 W halogen lamp and a 150 W projector (see **Figure S1** for a schematic representation). FR-PSI was alternatively photoaccumulated with far-red light by illumination with a 1 kW halogen lamp filtered by an IR filter and a long-pass 780 nm filter, yielding negligible transmitted light below 725 nm (see **Figure S1** for a schematic representation of the setup and transmittance profile of the combination of filters).

**Figure S1.**
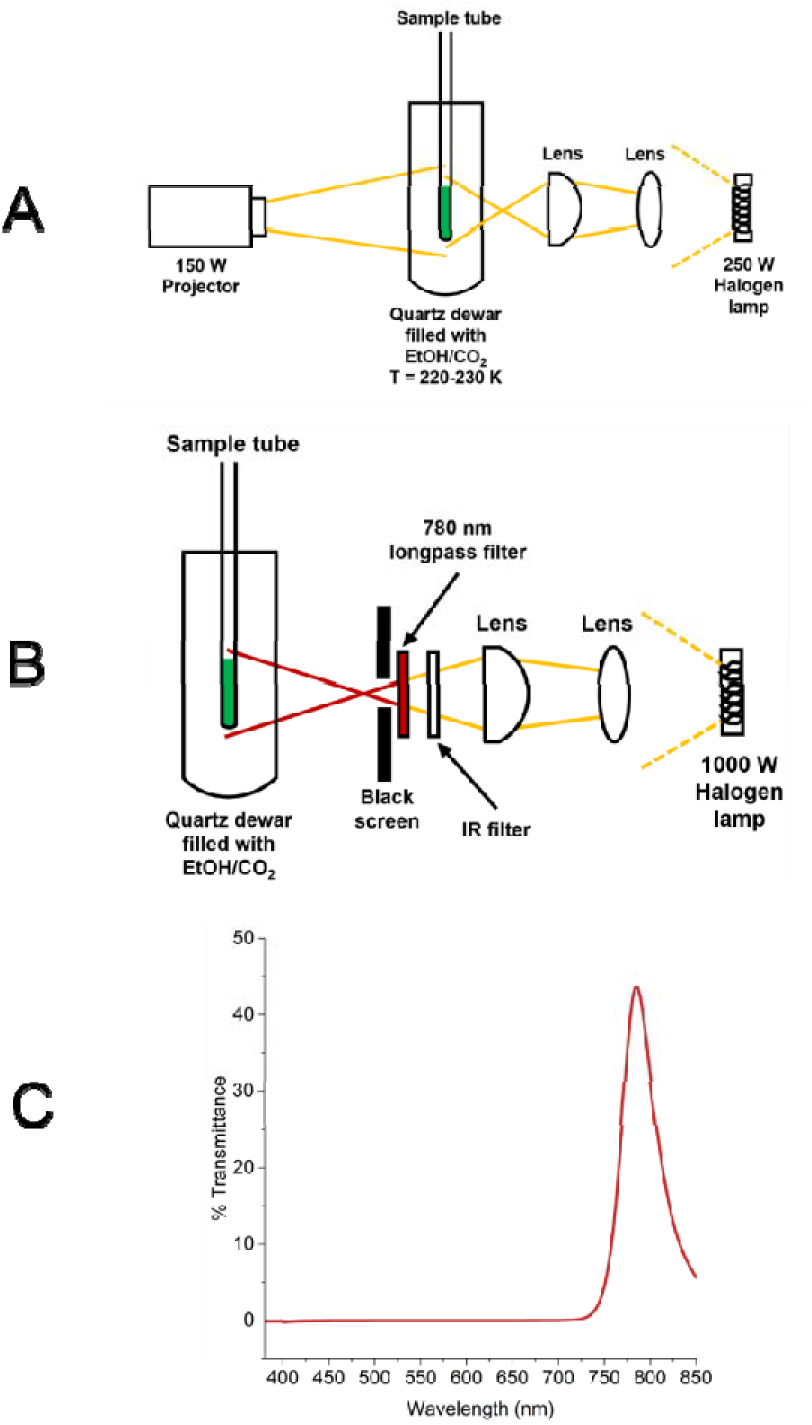
A: Experimental photoreduction setup with broad visible light. **B:** Experimental photoreduction setup with FR light. **C:** Transmittance profile of the combination of IR filter + 780 nm longpass filter used for FR light illumination.

#### Additional data analysis description: Procedure for the simulation of Q-band TR-EPR spectra

Q-band TR-EPR spectra were simulated starting by defining the orientations of the principal axes reference frames of all the involved tensors. The principal axes of the quinone ***g***-tensor for A_1A_ (*x̂_A_*, *ŷ_A_*, *ẑ_A_*) and A_1B_ (*x̂_B_*, *ŷ_B_*, *ẑ_B_*) were retrieved from the crystal structure of PSI (PDB: 1JB0)^3^, as they typically coincide with the symmetry properties of the quinone’s molecular structure. *x̂* axis was identified by the four aligned atoms composing the two C=O groups (direction obtained via singular value decomposition) and established in order to point away from P_700_. *ŷ* was found via singular value decomposition of the reference points obtained by calculation of the centres of mass between the carbon atoms C2-C3, C5-C10 and C7-C8 (atom numbering of the 1JB0 file) and pointing towards the position of the phytyl chain. *ẑ* was naturally fixed to be perpendicular to *x̂* and *ŷ*, and thus perpendicular to the quinone plane.

For the dipolar interaction, the dipolar principal axis *ẑ_d_* was aligned along the segment connecting the chlorophyll and quinone centres of spin distribution, pointing towards P_700_. For chlorophyll, the centre of spin distribution was estimated as the centre of mass between the four nitrogen atoms composing the chlorin ring of P_B_ (chlorophyll coordinated by PsaB and located on the B-branch side). For A_1A_ and A_1B_, their centres of spin distribution were obtained calculating the centres of mass of the two C=O groups, separately for both quinones. Due to the axial symmetry of the dipolar interaction, the principal axes *x̂_d_* and *ŷ_d_* can be set arbitrarily.

At variance with quinone ***g***-frames, the principal axes reference frame of chlorophyll radical ***g***-tensor (*x̂_Chl_*, *ŷ_Chl_*, *ẑ_Chl_*) does not follow the geometrical properties of the molecule.^4^ Therefore, it was identified differently: chlorophyll ***g***-frame was oriented in space so that the direction of the dipolar axis connecting A_1A_ and P_B_ is described by the set of polar angles (*θ*, *ϕ* = 39°, 62°) with respect to the chlorophyll ***g***-tensor itself. This procedure actually identifies an infinite family of chlorophyll ***g***-tensor reference frames, as whatever rotation around the direction of the just mentioned dipolar axis conserves the (*θ*, *ϕ* = 39°, 62°) polar coordinates. To obtain a unique chlorophyll ***g***-tensor, the reference frame was gradually rotated in order to obtain a set of polar angles (*θ*, *ϕ* = 90°, 54°) describing the orientation of the dipolar axis connecting A_1B_ and P_B_ with respect to the chlorophyll ***g***-tensor itself. In this way, a “double constraint” is imposed, which makes it possible to define a single orientation of the chlorophyll ***g***-tensor (without considering allowed symmetry transformations). The two couples of polar angles were chosen in agreement with literature data.^5^

All principal axes reference frames of all the involved tensors (visualized in **Figure 8** in the main text), that are actually defined with respect to the crystallographic reference frame of the pdb 1JB0 file, were inserted into the *eulang()* function of the EasySpin toolbox^6^ in order to obtain Euler angles describing the rotation of the crystallographic reference frame (chosen as general molecular reference frame) into the tensors’ reference frames, which is a required step for simulations with EasySpin.

All other simulation parameters, i.e. *g* principal al values, electronic couplings and linewidths are reported in **Table 3** in the main text.

#### Additional Results and Analysis

**Figure S2.**
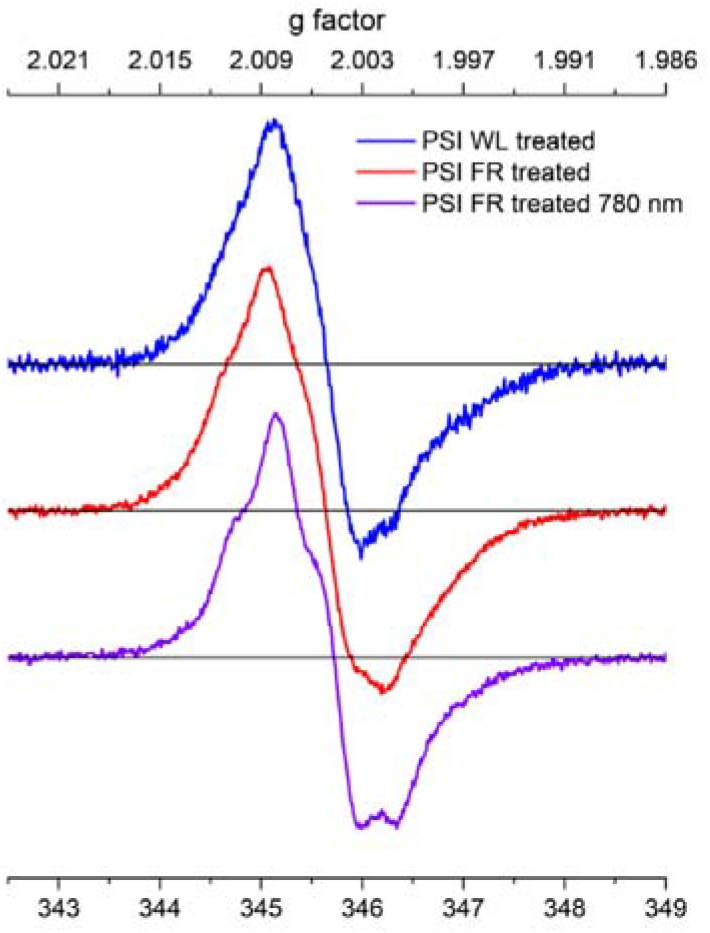
X-band CW-EPR spectra of quinone radicals produced in photoreduced WL-PSI (blue line), FR-PSI (red line) and FR-PSI under 780 nm filtered light (purple line). Experimental conditions: T = 80 K, modulation frequency = 40 kHz, modulation amplitude = 0.1 mT, MW power = 10 mW (41 dB), time constant = 20 ms, conversion time = 80 ms, magnetic field sweep = 7 mT (0.01 mT step), about 20 scans per spectrum. For direct comparison, the presented spectra are horizontally shifted through a common reference frequency of 9.7 GHz. MW cavity and cryostat as described in the main text.

**Figure S3.**
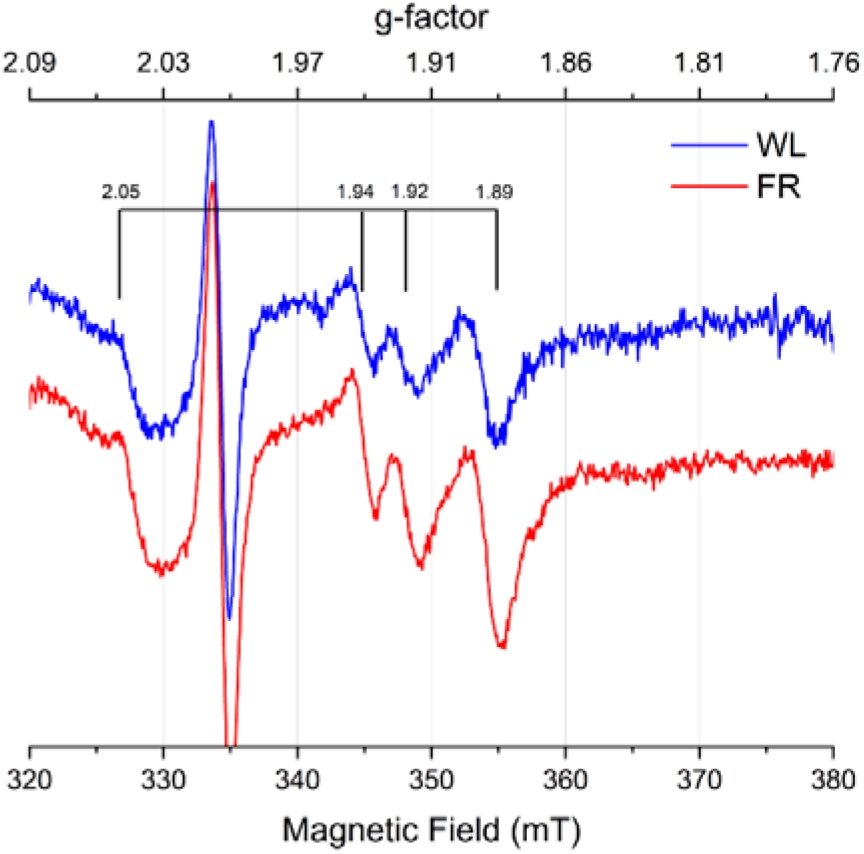
X-band CW-EPR spectra showing the typical F_A_/F_B_ interaction spectrum in photoreduced WL-PSI (blue line) and FR-PSI (red line). Experimental conditions: T = 9 K, modulation frequency = 100 kHz, modulation amplitude = 1.5 mT, MW power = 20 mW (10 dB), time constant = 10 ms, conversion time = 40 ms, magnetic field sweep = 60 mT (0.08 mT step), about 20 scans per spectrum. For these experiments, the MW cavity was a Bruker 4102ST/9901 critically coupled resonator working at 9.37 GHz and mounted on an ESR900 (Oxford) liquid helium cryostat.

**Figure S4.**
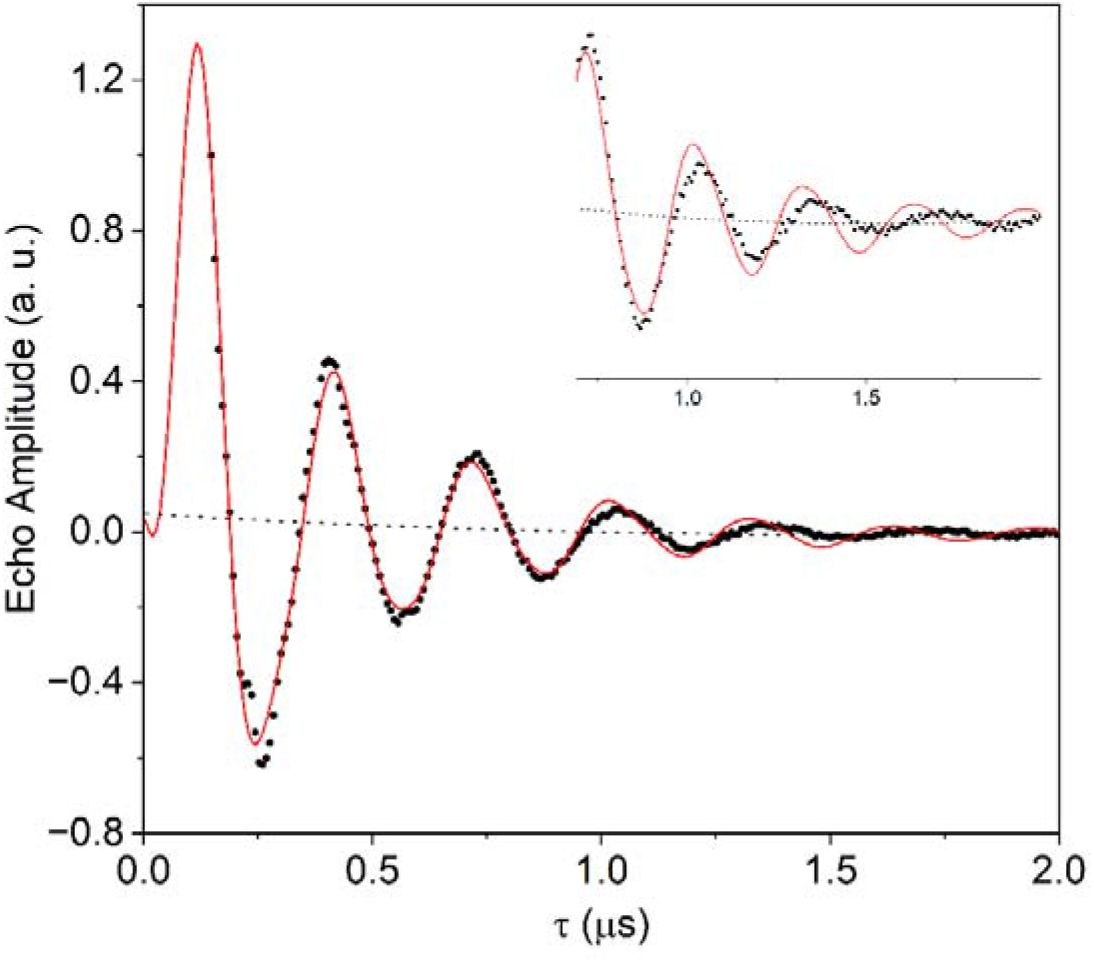
Fitting of the oop-ESEEM time trace (DAF = 0.3 ms) for dark adapted PSI-FR using a single modulation frequency component (instead of two as described in the main text). Experimental data: black dots; Simulation: red line. The failure of the fitting is highlighted by the inset, where predicted modulation beyond t = 1 ms does not correspond to the experimental trace

**Figure S5.**
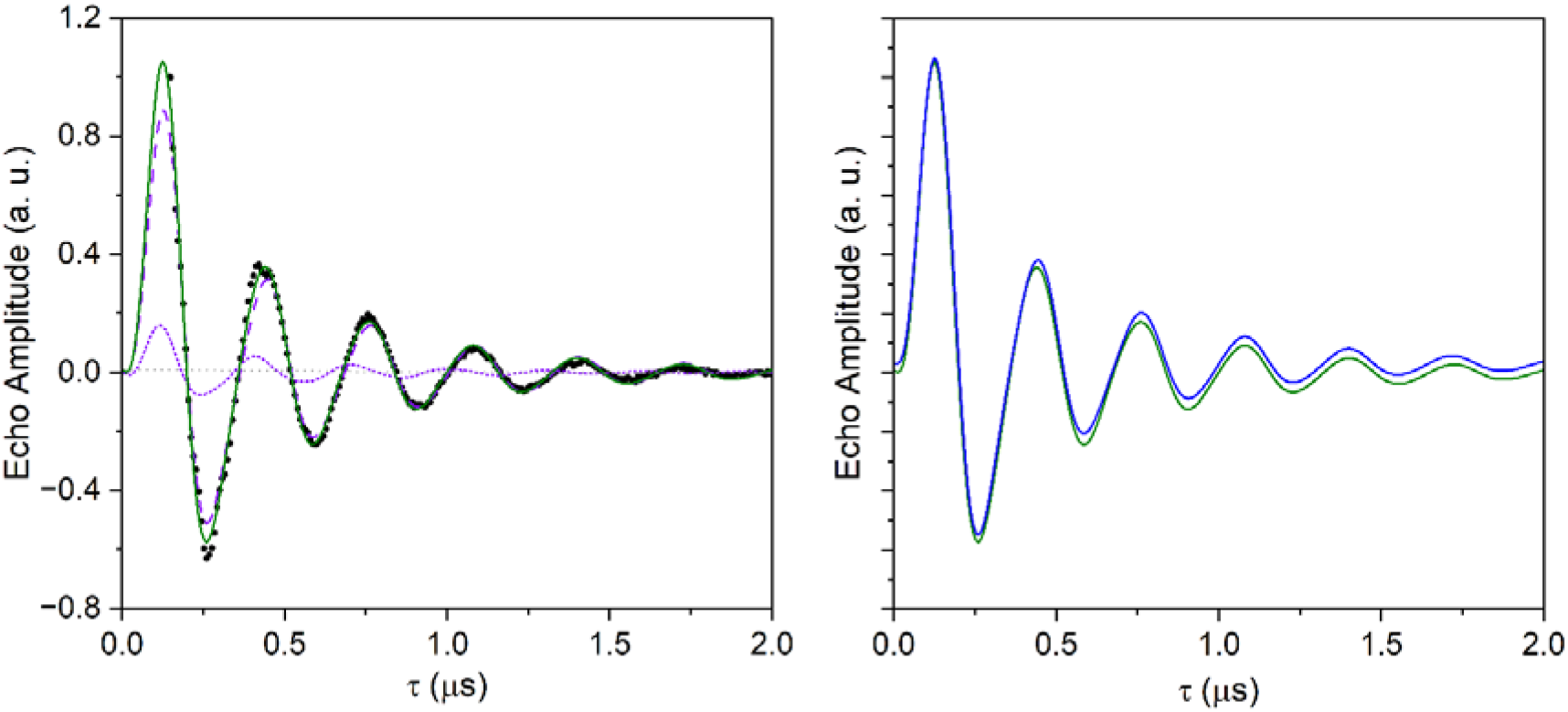
Fitting of the oop-ESEEM time trace (DAF = 0.3 ms) for dark adapted PSI-WL. **On the left**: experimental data (reported in black dots) fitted with a sum of two modulation frequencies, *i.e.* a lower modulation frequency accounting for [P_700_^+^A_1A_^−^] (in magenta dashed line), and a higher one accounting for [P_700_^+^A_1B_^−^] (in magenta dotted line), with relative amplitudes reflecting the ESE kinetic decay analysis. Total fitting shown as a green line. Baseline reported with dotted grey line. **On the right**: comparison of the fitting traces considering only a single frequency component (blue line, the same as reported in the Main Text, accounting for [P_700_^+^A_1A_^−^]) or two frequency components (green line). Traces are vertically shifted for clarity. No readily observable differences can be noted between the two fittings, in support of the fact that oop-ESEEM traces for WL-PSI (at any DAF) can be satisfactorily reconstructed by considering only [P_700_^+^A_1A_^−^] contribution.

**Figure S6.**
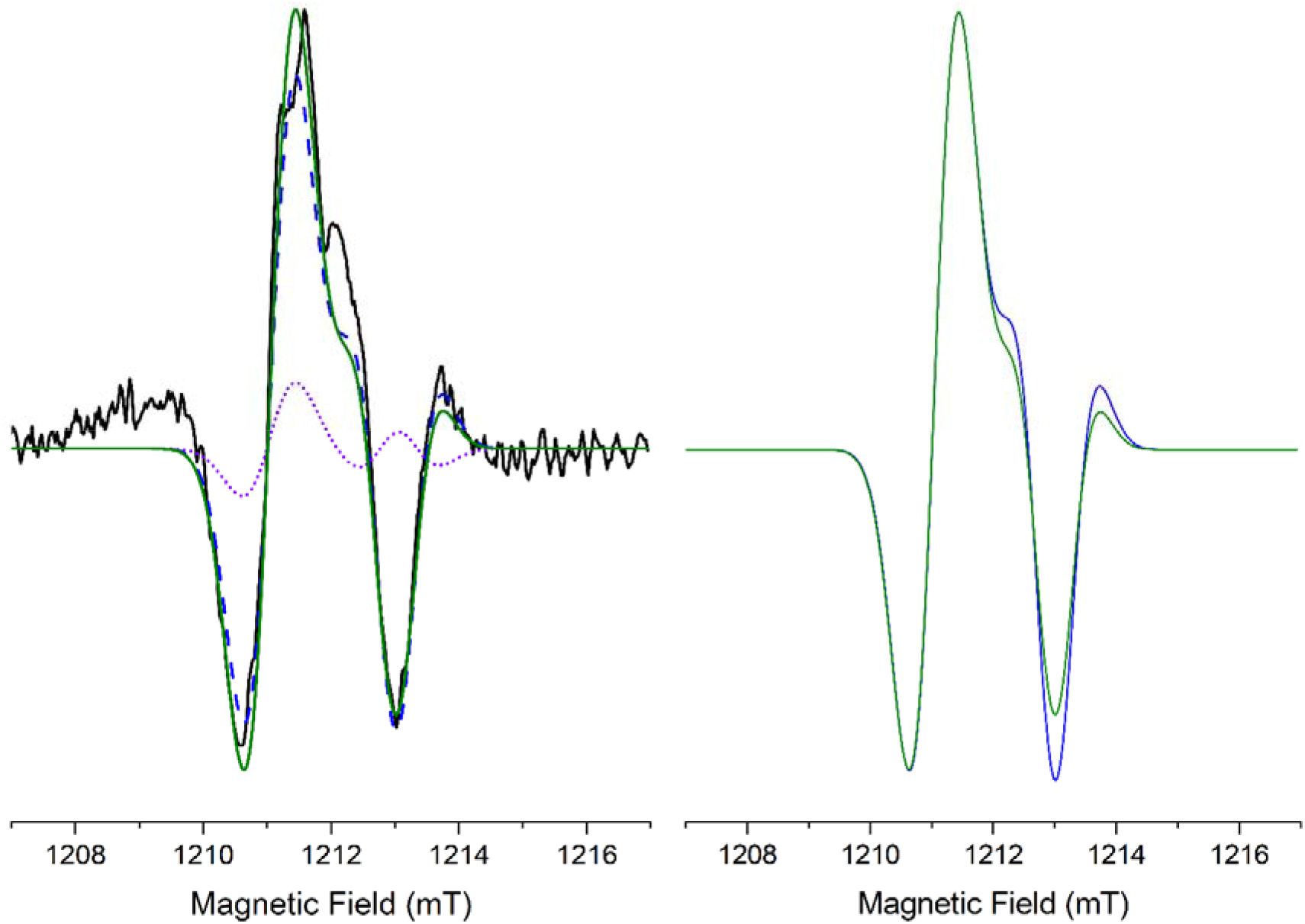
Simulation of the Q-band TR-EPR spectrum for dark adapted PSI-WL. **On the left**: experimental spectrum (reported in black line) simulated with a weighted sum of two components accounting for [P_700_^+^A_1A_^−^] (dashed blue line) and [P_700_^+^A_1B_^−^] (purple dotted line), with relative amplitudes reflecting the ESE kinetic decay analysis. Total simulation shown as a green line. **On the right**: comparison of the simulation traces considering only a single component (blue line, the same as reported in the Main Text, accounting for [P_700_^+^A_1A_^−^]) or two components (green line). Differences are evident in the high field portion of the spectrum. The simulation with two components better describes the experimental negative feature at 1213 mT compared to the simulation with just a single component, but the reconstruction is worse considering the positive feature at 1213.5 mT. These observations conclude that no net gain in the simulation quality is achieved when WL-PSI TR-EPR spectrum is simulated with two components, justifying its satisfactory simulation with a single term accounting for [P_700_^+^A_1A_^−^].

